# Brain-optimized neural networks learn non-hierarchical models of representation in human visual cortex

**DOI:** 10.1101/2022.01.21.477293

**Authors:** Ghislain St-Yves, Emily J. Allen, Yihan Wu, Kendrick Kay, Thomas Naselaris

## Abstract

Deep neural networks (DNNs) trained to perform visual tasks learn representations that align with the hierarchy of visual areas in the primate brain. This finding has been taken to imply that the primate visual system forms representations by passing them through a hierarchical sequence of brain areas, just as DNNs form representations by passing them through a hierarchical sequence of layers. To test the validity of this assumption, we optimized DNNs not to perform visual tasks but to directly predict brain activity in human visual areas V1–V4. Using a massive sampling of human brain activity, we constructed brain-optimized networks that predict brain activity even more accurately than task-optimized networks. We show that brain-optimized networks can learn representations that diverge from those formed in a strict hierarchy. Brain-optimized networks do not need to align representations in V1–V4 with layer depth; moreover, they are able to accurately model anterior brain areas (e.g., V4) without computing intermediary representations associated with posterior brain areas (e.g., V1). Our results challenge the view that human visual areas V1–V4 act—like the early layers of a DNN—as a serial pre-processing sequence for higher areas, and suggest they may subserve their own independent functions.

## 1 Introduction

Primate visual cortex contains dozens of functionally distinct areas. Understanding how these areas relate to one another and identifying the function or functions of the different representations in these areas has been a driving pre-occupation of visual neuroscience for decades [Carandini et al., 2005, Roe et al., 2012, DiCarlo and Cox, 2007].

Studies of the so-called “early” visual areas V1, V2, V3 and V4 have played an important role in understanding the diversity of functions supported by the many brain areas in primate visual cortex. Each of V1–V4 represents a complete map of visual space whose boundaries are identifiable on cortical surface maps [Hansen et al., 2007, Wandell and Winawer, 2011]. Importantly, visual representations at a given location in visual space vary dramatically across areas [Kobatake and Tanaka, 1994, Grill-Spector and Malach, 2004]. V1–V4 thus offer an opportunity to understand the principles by which visual representations are transformed when they cross a boundary between brain areas.

Although we currently lack a complete characterization of the representations in V1–V4, the belief that they are part of a hierarchy of visual processing is deeply entrenched in both neuroscience and AI [Hubel and Wiesel, 1962, Riesenhuber and Poggio, 2000, LeCun et al.,1989, Krizhevsky et al., 2012, Felleman and Van Essen, 1991, Richards et al., 2019, Himberger et al., 2018, Yamins and DiCarlo, 2016]. According to this belief, representations in V1 are the least complex in the hierarchy; they are passed upward through V2 and V3 to V4 and beyond, where representations become more complex.

Belief in this hierarchy of representations is founded on well-established evidence for an anatomical hierarchy defined by the laminar distributions of reciprocal connections between pairs of brain areas [Felleman and Van Essen, 1991], the evidence for a hierarchy of spatial resolution defined by the monotonic increase in receptive field sizes [Dumoulin and Wandell,2008, Kay et al., 2013b] and monotonic decrease in preferred spatial frequencies [Henriksson et al., 2008], and the evidence for a temporal hierarchy defined by the increase in neural onset latencies [Schmolesky et al., 1998] across V1–V4.

Belief in a hierarchy of representations has had a profound impact on the design of artificial systems that solve visual tasks [Lecun et al., 2015]. Specifically, the architecture of convolutional deep neural networks (DNNs) specifies a hierarchy of visual maps [Fukushima, 1988]. The success of DNNs at solving hard vision tasks [Krizhevsky et al., 2012] has been interpreted as providing evidence for a hierarchy of representations in the brain [Lindsay, 2021]. Even stronger support for a hierarchy of representations in the brain comes from the well-established finding that “task-optimized” DNNs trained on supervised [Yamins et al., 2014, Khaligh-Razavi and Kriegeskorte, 2014, Güçlü and van Gerven, 2015, Cichy et al., 2016, Eickenberg et al.,2017], and unsupervised [Zhuang et al., 2021] computer vision tasks align their layers to the presumed hierarchy of visual brain areas.

Despite the strong evidence appearing to support the idea that the primate visual system implements a “hierarchy of representations”, the phrase remains a verbal summary of a diverse set of experimental observations [Hilgetag and Goulas, 2020] and has not yet, to our knowledge, been distilled into a set of key properties that visual representations must satisfy in order to qualify as hierarchical. To do so, we need to articulate these key properties and develop falsifiable tests for each. Given that our evolving understanding of representational organization in the brain has had a significant causal impact on the development of AI [Macpherson et al.,2021, Hassabis et al., 2017], a direct test for hierarchical representation in the primate visual system could yield important insights for both our understanding of the brain and the design of intelligent machines.

In this work, we define hierarchical representations as those that are built up through the iterative application of a fixed transformation [Yamins and DiCarlo, 2016]. If V1–V4 conform to this definition then each brain area can be viewed as a layer in a single DNN. Independently falsifiable tests for hierarchical representation in V1–V4 follow naturally: First, predictive encoding models of V1–V4 [Naselaris et al., 2011] should perform better or worse to the extent that they are more or less consistent with this definition of hierarchy. Second, accurate DNN models of representation in anterior brain areas (e.g., V4) should demand more depth than models of posterior brain areas (e.g., V1). Third, since representations in a hierarchy are computed in sequence, accurate models of representations in anterior brain areas should always *entail* (require the construction of) representations encoded in posterior areas. If V1–V4 do not meet some or all of these criteria, we must conclude that the representations they encode are non-hierarchical.

To test if these criteria of hierarchical representations are met by V1–V4, we train neural networks to predict human brain activity [Seeliger et al., 2021, Cadena et al., 2019, Prenger et al., 2004, Antolík et al., 2016, Batty et al., 2016, Klindt et al., 2017, McIntosh et al., 2016,Kindel et al., 2019, Zhang et al., 2019] in a massive sampling of responses to hundreds of thousands of presentations of natural scenes [Allen et al., 2022]. By contrasting such “brain-optimized networks” to task-optimized networks, we are able to discriminate representations that may be essential for solving computer vision tasks but play no essential role in modeling the brain. By analyzing the accuracy with which these diverse networks predict brain activity, the coupling between the layers of these networks and brain areas V1–V4, and the ability of representations optimized for one brain area to generalize to others, we are able to conduct novel and sensitive tests for the existence of hierarchical representation in human visual cortex.

## 2 Results

### 2.1 Three tests for hierarchical representation

We consider deep neural network models of representation in V1–V4. In a DNN, representations are constructed via compositions *e*_l_(*x*) of a transformation *η_θ_l__*(*x*):

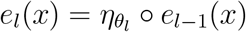

where *x* is a visual stimulus, subscripts *l* index particular values of the adjustable parameters *θ*, and *e*_0_(*x*) = *x*. We denote the complete set of representations expressed by the network as *ϕ* ≡ (*e*_1_, …, *e_L_*), where *L* is number of layers in the DNN.

We define two general properties that DNN-based models of representation in V1–V4 must satisfy in order to be considered “hierarchical”. The first property, “ordering”, is a generalization of the finding that task-optimized DNNs order V1–V4 with respect to layer depth. Let 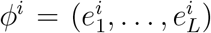 and 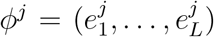 be representations we use to predict brain activity in voxel group *V_i_* and *V_j_*, respectively. Here, *i,j* ∈ (1, 2, 3, 4) correspond to the visual area of the same index. We assume that all *ϕ*’s have the same depth *L* and utilize transformations of the same form *η*, but may or may not correspond to identical DNNs. We say that *ϕ^i^* and *ϕ^j^* are ordered with respect to *V_i_* and *V_j_* for a partitioning at layer l if representations below 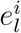 contribute more strongly (relative to other layers) to predicting brain activity in *V_i_* than representations below 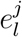 contribute to *V_j_* whenever *i* < *j*.

The second property, “entailment”, is a generalization of the dependence of higher layers on lower layers in a DNN. In a DNN, representations are computed in sequence. This means representations in lower layers must be computed in order to compute representations in higher layers. In our work, entailment is a generalization of this hierarchical dependence that can be applied to distinct sets of representations *ϕ^i^* and *ϕ^j^*. Specifically, we say that *ϕ^i^* and *ϕ^j^* show entailment with respect to *V_i_* and *V_j_* if the representations in *ϕ^i^* that contribute to predicting brain activity in *V_i_* are a subset of the representations in *ϕ^j^*, whenever *i* < *j*.

When representations show both ordering and entailment, we say they are hierarchical with respect to V1–V4 (Fig. 1a); when they show neither property we say they are non-hierarchical with respect to V1–V4 (Fig. 1b).

**Figure 1:**
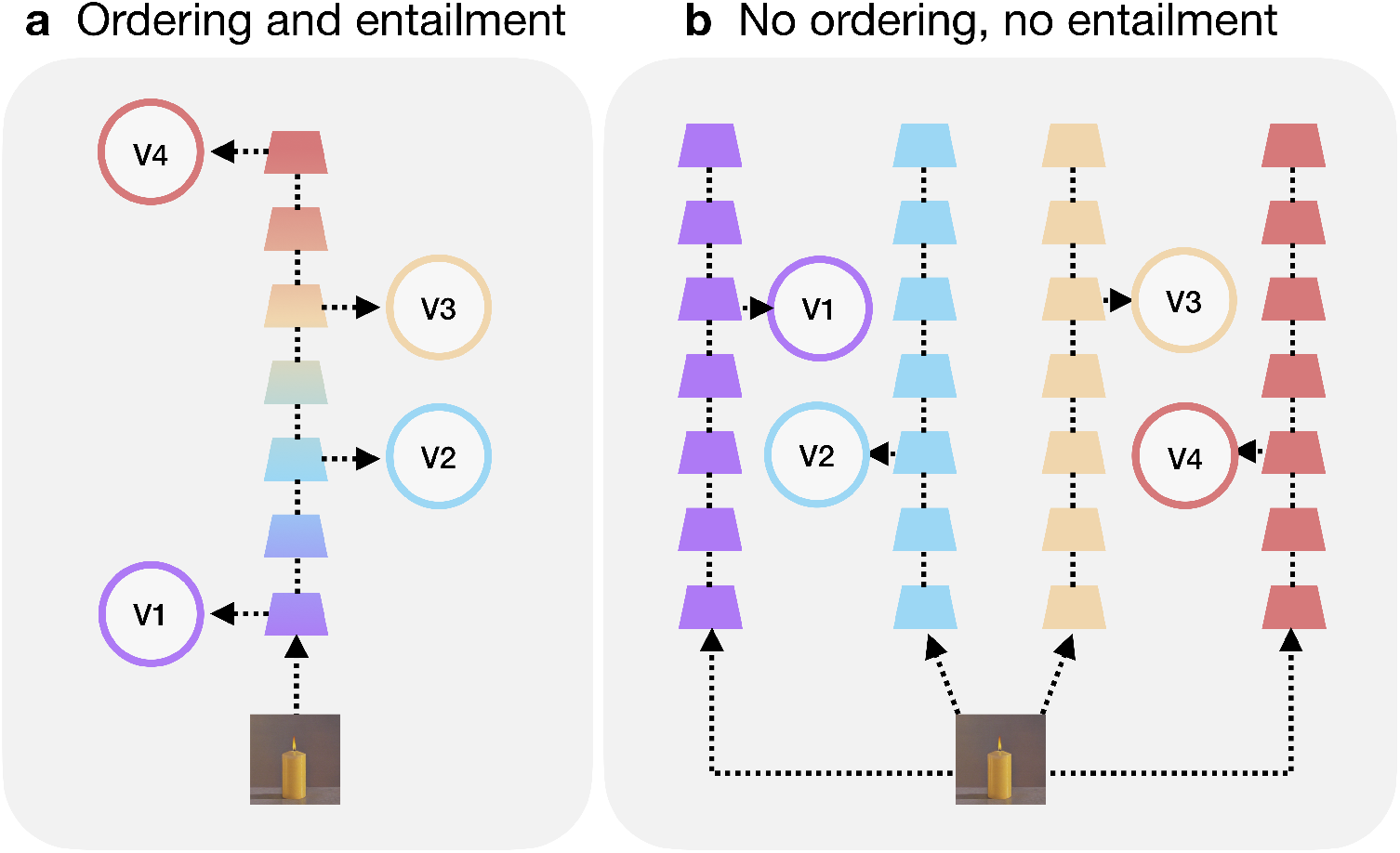
Examples of hierarchical and non-hierarchical representations. (**a**) An example of hierarchical representation in V1–V4 (circles). The depth of layers that contribute (dashed arrows) to predicting brain activity are aligned to V1–V4 (ordering). Since representations in these layers must be computed in sequence, we would infer that representations in V4 entail representations in V3, and so on. (**b**) An example of non-hierarchical representation in V1–V4. The depth of layers that contribute to predicting brain activity are roughly the same and do not align with V1–V4 (no ordering). Representations in these contributing layers can be computed in parallel, so in this example there is no evidence that representations in any area entail representations in any other area. In these simplified illustrations only a single layer contributes to predicting activity in each brain area; note, however, that is not a requirement. In our models all layers may contribute to predicting activity in any brain area.

With these definitions in place, we can construct tests for properties that we expect a hierarchical representation to satisfy by analyzing DNNs that are optimized to predict activity in V1–V4.

First, if V1–V4 encode hierarchical representations, then models that are hierarchical with respect to V1–V4 should predict brain activity more accurately than those that are not. This implies that a network training strategy that facilitates the learning of hierarchical representations should be a useful inductive bias [Goyal and Bengio, 2020]. Specifically, we expect that a single DNN trained jointly on V1–V4 should yield more accurate predictions of brain activity than four distinct DNNs trained independently on each brain area. To see why this is a reasonable expectation, assume we want to construct a DNN to model V4. If V1–V4 are hierarchical, the DNN will predict activity in V4 most accurately if it entails representations that accurately model V1–V3. By optimizing the DNN jointly, we supply the learning algorithm with samples of brain activity that encode these lower-level representations. In contrast, if we train the DNN on activity sampled from V4 only, we deprive the learning algorithm of informative data and should therefore expect worse performance from the DNN once training is complete.

If V1–V4 encode hierarchical representations, then DNNs optimized to predict activity in V1–V4 should be ordered. Importantly, ordering should hold regardless of whether we train a single DNN on all areas or train four DNNs independently on each. By assessing ordering in independently trained DNNs, we can determine if alignment between brain areas and layers is simply an optimal arrangement for task-optimized networks specifically, or if it indicates that the diversity of representations across V1–V4 can only be accurately modeled by varying compositional depth.

Finally, if V1–V4 conform to our definition of a hierarchy, representations in anterior areas should entail representations in posterior areas. This implies, for instance, that if we train a DNN to predict V4 brain activity alone, we should also be able to use it to predict brain activity in V1–V3 as accurately as a DNN trained to model those more posterior areas independently. On the other hand, a DNN trained to predict V1 activity alone should not be expected to predict activity in V2–V4 as well as DNNs trained to model those more anterior areas independently. A network, or set of networks, that do not exhibit this hierarchical prediction asymmetry should not predict brain activity as accurately as one that does.

The tests we propose for identifying hierarchical representation in V1–V4 presume an ability to optimize neural networks to predict brain activity. In what follows, we first confirm that brain-optimized networks, when trained on sufficient amounts of data, can be made to yield state-of-the-art prediction accuracy in V1–V4, even for out-of-sample testing stimuli. We then use brain-optimized network models to perform the tests described above.

### 2.2 Encoding models based on brain-optimized networks yield accurate predictions of brain activity to natural and artificial stimuli

In encoding models based upon deep neural networks, the DNN acts a nonlinear feature extractor, and the activities of units in each feature map of the DNN are transformed into predicted brain activity via a linear read-out head (Fig. 2; see St-Yves and Naselaris [2018] as well as Allen et al. [2022]). For each voxel in the target dataset, the read-out head specifies a spatial receptive field and an array of feature weights that model the region of visual space and the nonlinear features that are represented by brain activity measured in the voxel. In our work the read-out head for each voxel always samples from all layers throughout the depth of the network. Thus, we give each layer in the feature-extractor network the chance to contribute to predicting brain activity (Fig. 2a,b).

**Figure 2:**
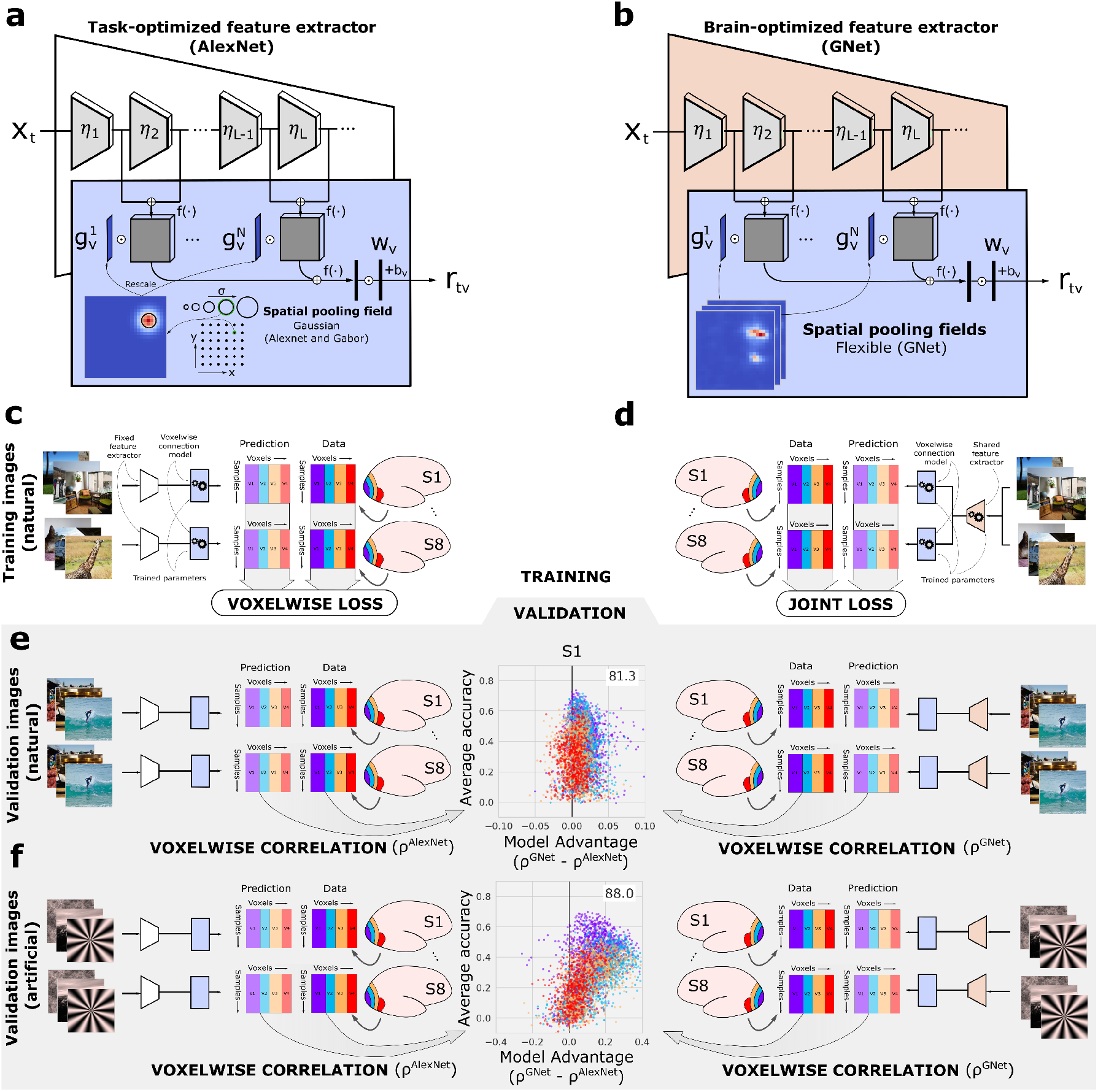
Training and validation of task-optimized and brain-optimized networks. (**a**) An encoding model based on a task-optimized deep neural network (AlexNet; white trapezoid). Multiple convolutional layers (*η_i_*) convert the input stimulus on trial *t, X_t_*, into feature maps. (**b**) A read-out head (blue trapezoid) transforms network activations into predicted brain activity (*r_tv_*, where *v* indexes a single voxel). The read-out head consists of Gaussian spatial pooling fields (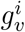; example at lower left) with position and size selected from a fixed grid of candidates (lower right). The pooling field and each feature map are multiplied pixel-wise and then summed, reducing each feature map to a single feature value. The array of feature values across all maps (left vertical rectangle) are weighted by an array of feature weights (*w_v_*) and then summed (with a bias term *b_v_*) to yield predicted brain activity. Compressive point nonlinearities (*f*(·)) are applied at several processing stages. (**b**) A similar architecture is used for the encoding model based on the brain-optimized network (GNet; orange trapezoid), although the “flexible” spatial pooling fields used in the read-out head may be non-Gaussian (example at lower left). (**c**) Training for the AlexNet-based encoding model. For each voxel, only readout head parameters (*g_v_,w_v_, b_v_*) are optimized (gears) for visual areas V1–V4 (colored rectangles) in each subjects’ brain (S1, … S8). The “voxelwise loss” (squared difference between predictions and measured activity data) is evaluated independently for each voxel. (**d**) Training for the GNet-based encoding model. Both the read-out head and GNet parameters are optimized jointly (“joint loss”) for all voxels, subjects and brain areas. (**e**) Prediction accuracy is evaluated for each voxel by correlating predicted brain activity with measured brain activity (*ρ*^AlexNet^, *ρ*^GNet^) across a set of held-out validation trials with natural scenes. We plot the average accuracy (y-axis of central plot) vs. the difference in accuracy (x-axis) for each voxel (dots; color indicates visual area). For this example subject S1 the GNet-based encoding model predicts responses to natural scenes most accurately for 81.3% of voxels in V1–V4. (**f**) In this example, the GNet-based encoding models predicts responses to artificial stimuli more accurately for 88% of voxels in V1–V4.

In order to perform tests of hierarchical representation, we constructed encoding models in which the feature extractor is a brain-optimized neural network (GNet, Fig. 2b). In the GNet encoding models, we minimized error on predicted brain activity by using stochastic gradient descent to learn all free parameters of the DNN feature extractor simultaneously with the free parameters of the read-out heads (Fig. 2d). We compared the prediction accuracy of these GNet-based encoding models to models based upon a task-optimized neural network (AlexNet; Krizhevsky et al. [2012]) that was pre-trained to discriminate object categories. For the AlexNet encoding model (Fig. 2a) the network parameters were frozen during training and only the free parameters of the read-out heads were optimized (Fig. 2c).

To optimize the parameters of each type of encoding model (Fig. 2c,d) we used the Natural Scenes Dataset [Allen et al., 2022], a massive sampling of blood-oxygenation-level-depedendent (BOLD) activity in eight subjects using ultra-high field fMRI (7T, 1.8-mm resolution). Subjects each viewed 9,000-10,000 natural scenes (sampled from the Microsoft Common Objects in Context database [Lin et al., 2015]) presented (3-s exposure) repeatedly (three times typically), yielding 22K - 30K trials for individual subjects and a total of 213K trials across subjects.

Voxels were assigned to areas V1–V4 on the basis of an independent retinotopic mapping experiment [Allen et al., 2022].

To validate and compare encoding models, after training we assessed the prediction accuracy of the models for each voxel by correlating predicted activity with measured activity in response to the shared images that were shown to all eight subjects during the experiment but were not used for model training (Fig. 2e). For each subject, the brain-optimized GNet encoding model (trained jointly) predicted brain activity in V1–V4 more accurately than the AlexNet encoding model for more than 68% of all voxels in V1–V4 (Fig. 3a). Averaged across subjects, the “win percentage” for the GNet model was significantly greater than expected by chance (80%win, *p* < 10^−6^, two-sided t-test). Interestingly, when model prediction accuracy was computed for out-of-sample classes of artificial stimuli (e.g., gratings, contrast-modulated scenes, various types of noise, Fig. 2f and Fig. S1), the prediction accuracy of the GNet model was greater than the AlexNet model for more than 76% of the voxels in all subjects (Fig. 3b). Although the size of the difference in prediction accuracy underlying these win percentages varied within and across areas (Fig. 3c), across subjects the average win percentage of the GNet model for artificial stimuli was significantly greater than for natural stimuli (80% win, *p* < 10^−5^, two-sided t-test). The win percentage was not significantly different across these two stimulus conditions for any single brain area except V4, where we observed an average improvement across subjects from 62%to 74% (*p* < 0.01, two-sided paired t-test) (Fig. 3d).

**Figure 3:**
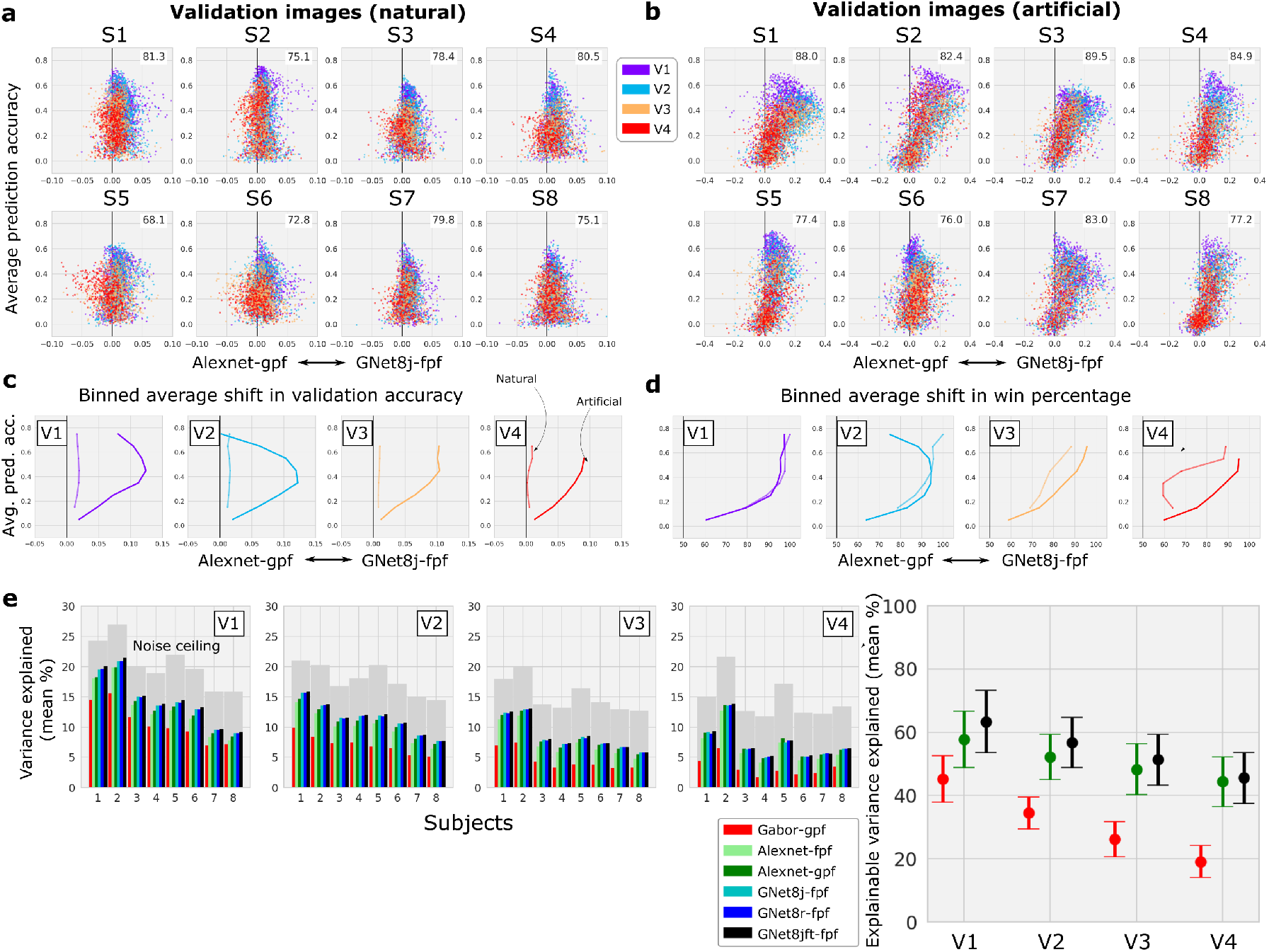
Comparison of cross-validated prediction accuracy for encoding models based on task- and brain-optimized deep neural networks. (**a, b**) Accuracy / advantage plots for all subjects and brain areas for natural validation stimuli (a) and artificial stimuli (b). Format as in Figure 1. (**c**) Median difference in model prediction accuracy (x-axis) for different levels of average prediction accuracy (y-axis) and for natural (thin curve) and artificial (thick curve) stimuli. (**d**) Difference in the percentage of voxels for which the the GNet-based encoding model explains more signal variance than AlexNet-based encoding model (“win percentage”; x-axis). (**e**) Signal variance (%; y axes) in brain activity explained by varieties of network-based encoding models (colored bars; gray bars indicate theoretical upper limit) for all each subjects (x-axis) in visual areas V1–V4. Inset: Average (across subjects) percent of explainable variance explained by a subset of the models. See Supplementary Table 1 for descriptions of all model acronyms.

These results demonstrate that the GNet model outperforms the AlexNet model for a majority of voxels in V1–V4. We also quantified the prediction accuracy of multiple model types with respect to the best performance that any model might achieve. To do this, we estimated the percentage of variance in brain activity that can be explained by variation in the stimulus (by computing noise ceilings as in Allen et al. [2022], gray bars in Figure 3e), as well as the percentage of variance explained by the outputs of each model (colored bars in Figure 3e). For every subject and brain area (with the exception of S4, V4), the mean (across all voxels) percentage of total variance explained by GNet models is larger than or equal to the mean percentage of variance explained by the AlexNet models and a control Gabor wavelet model. Improvements in prediction accuracy achieved by extensive fine-tuning of the model parameters (see Methods) suggests that the limit of prediction accuracy for GNet models has not yet been reached (“GNet8jft-fpf”, Fig. 3e, right). However, across all subjects and brain areas, GNet encoding models can account for at most 78% of the explainable variance, and as little as 37% (on average over a ROI, for voxels with at least 5% of explainable variance). This means that currently, even the best models leave much to be explained. Interestingly, this limitation does not seem to be unique to models trained on BOLD activity. For most subjects, the amount of explainable variance in V1 that is explained by GNet is similar to the amount explained by similar brain-optimized DNN-based models of single-unit activity in V1 of the primate brain [Cadena et al., 2019].

To further demonstrate the adequacy of brain-optimized models, we analyzed pooling fields (Fig. 4a) for voxels in V1–V4, recovering expected visual field coverage (Fig. 4b) and size-eccentricity relationships (Fig. 4c,d; [Kay et al., 2013b]). We conclude that our encoding models based upon brain-optimized networks are as trustworthy a tool for testing hypotheses about representation as any currently existing network-based encoding model.

**Figure 4:**
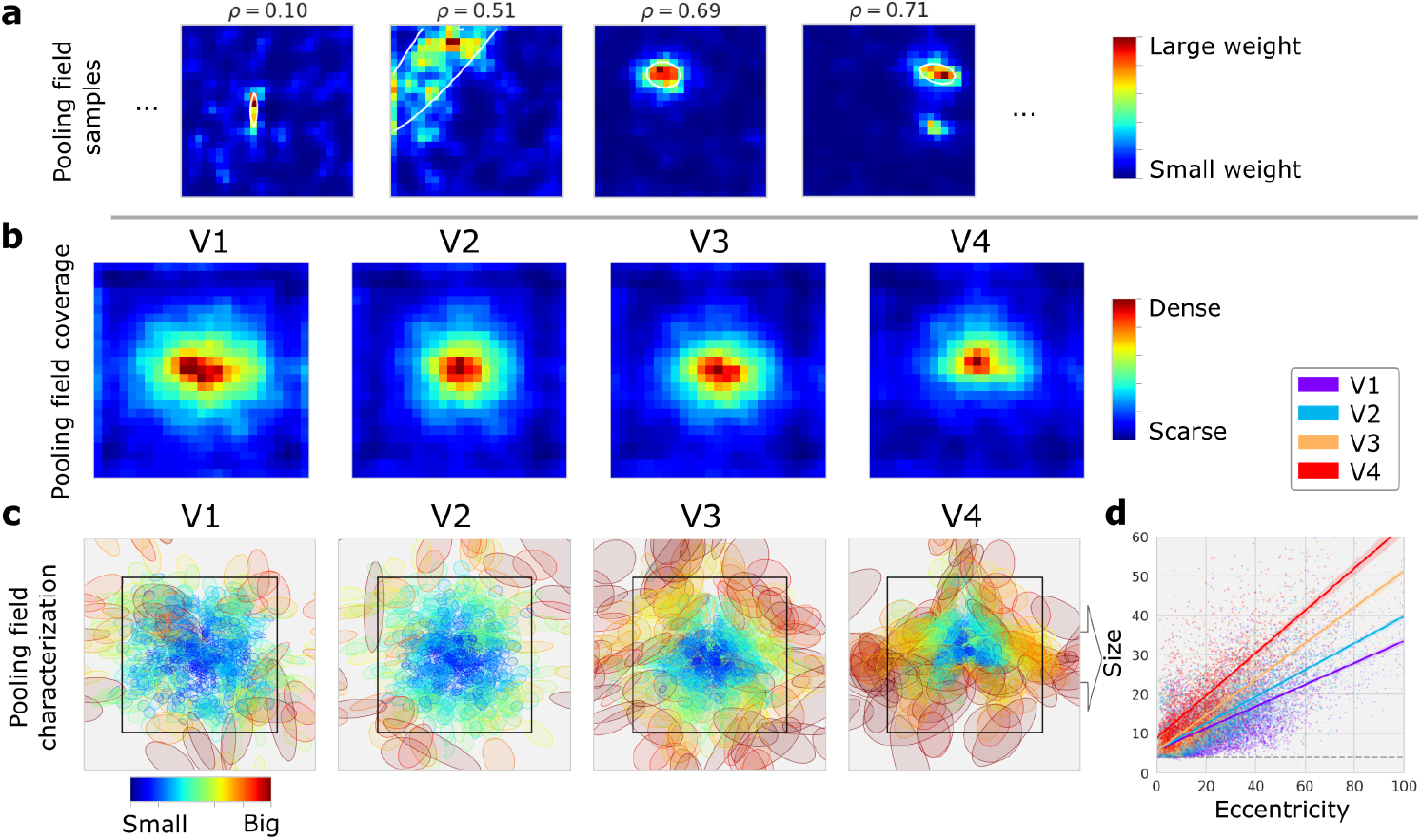
Encoding models based on brain-optimized networks recover known retinotopic organization. (**a**) Examples of spatial pooling fields (colormap indicates strength of predicted activation) and best-fitting Gaussian profile (with ellipsoids) for individual voxels with significantly accurate (*ρ* > 0.055, *ρ* < 0.01) encoding models. (**b**) Mean of spatial pooling fields for all voxels and subjects in V1–V4. (**c**) Best-fitting Gaussian profiles for spatial poolings of all voxels and subjects. The profiles were used to visualize size-eccentricity relationships for all visual areas. (**d**) Linear fits to relationship between pooling field size (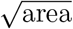 of the one std. dev. elliptical Gaussian profile in (c)) and eccentricity for all voxels in V1–V4. Length units are expressed in percent of stimulus span (i.e. 100% ≡ 8.4°, black bounding box in (c)).

### 2.3 Representational hierarchy is not an effective inductive bias for constructing predictive encoding models

To test the effectiveness of hierarchy as inductive bias, we compared a single GNet encoding model trained jointly on V1–V4 (GNet8j, where “8” indicates joint training for all 8 subjects, and “j” indicates joint training for all four brain areas, Fig. 5a) to four separate GNet encoding models trained independently to predict V1–V4 (GNet8r, where “r” stands for “ROI-wise”, Fig. 5b). The win percentages for GNet8j and GNetr were close to parity in V1–V3, although in V4 win percentage for GNet8r was 68%. Across several model variations, the win percentage of the GNet8j model was at best equal to the win percentage for the GNet8r model. This indicates that the jointly-trained GNet model was not more accurate than the independently trained GNet models (Fig. 5c). In fact, the independently trained models yielded better prediction accuracy than the jointly trained model, though the advantage is relatively small. Thus, a training strategy that favored discovery of hierarchical representations did not result in a relative increase in prediction accuracy over a training strategy that did not.

**Figure 5:**
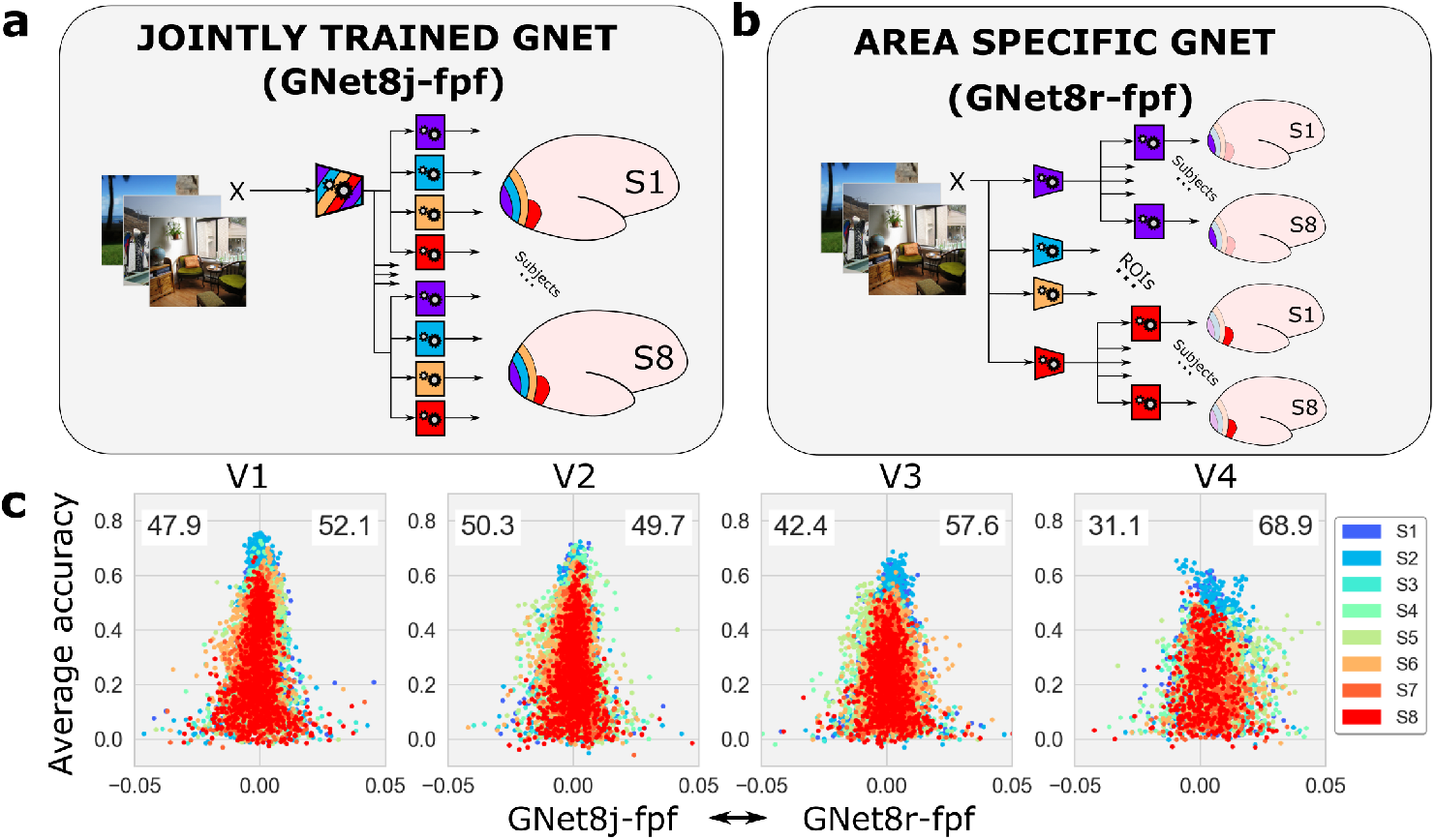
Comparison of brain-optimized DNNs trained jointly or independently on activity in V1–V4. (**a**) Training and architecture for the jointly-trained (GNet8j-fpf) variant of the GNet-based encoding model. (**b**) The independently-trained variant (GNet8r-fpf). A separate GNet feature extractor (trapezoids) is trained (gears) for each brain area (V1–V4). In both cases, read-out heads (squares) with flexible pooling fields (fpf) are optimized as well. (**c**) Advantage / accuracy plots comparing prediction accuracy of encoding models based on the jointly-trained (left of vertical line in each panel) and independently-trained (right of vertical line) GNets for each subject (colors).

### 2.4 Alignment between layers and V1–V4 does not indicate a requirement for compositional depth

Although GNet8r was not trained in a way that facilitated learning representations with ordering and entailment, it may have learned them anyway. To identify networks that order V1–V4, we tested the contribution of layers in the bottom half of the various feature-extracting DNNs to explaining variance in brain activity (Fig. 6, bold curves). We define the “specific” contribution of any set of layers as the prediction accuracy obtained when generating predictions with that set of layers alone (i.e., by masking out all other layers; Fig. 6b,c, blue curves). We define the “unique” contribution of any set of layers as the prediction accuracy of the full model, minus the specific contribution of all other layers (Fig. 6c, magenta curves). For two variants of the AlexNet encoding models (“AlexNet-gpf” and “AlexNet-fpf”), and for the jointly trained GNet encoding model (“GNet8j-fpf”), the unique contribution of lower layers declined monotonically from V1–V4 from roughly 60% to roughly 30%. Concurrently, the unique variance for the top layers (indicated by the distance from the top of the y-axis to the blue curves in Figure 6c) increased monotonically from V1–V4. Thus, the DNNs in these models do indeed induce an ordering with respect to V1–V4, as expected from previous results [Gücçlü and van Gerven,2015].

**Figure 6:**
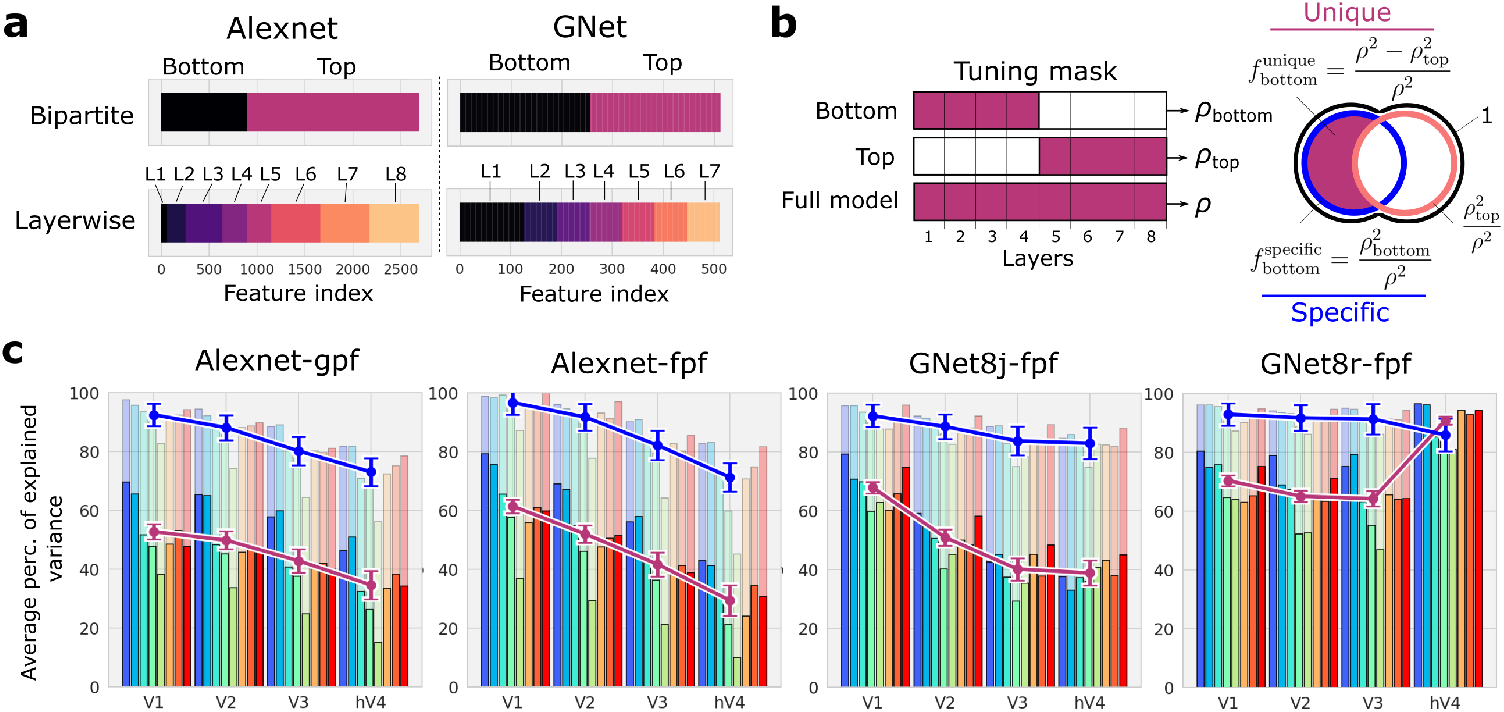
Alignment between layers and brain areas in task-optimized and brain-optimized networks. (**a**) Partitioning of layers (L1, L2,…) into bottom and top halves for AlexNet and GNet. (**b**) Partitioning of model output variance for bottom and top halves of the networks. Model predictions are generated using either the bottom or top layers alone, and the prediction accuracy is calculated (*ρ*_bottom_, *ρ*_top_). These quantities are used to calculate the “specific” 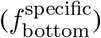 and “unique” 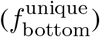 fractions of variance explained by the bottom layers. Both measure the contribution of the bottom layers to predicting activity in each brain area. (**c**) Specific (light bars, blue curve) and unique variance (solid bars, magenta curve) explained by the bottom layers for each brain area (x-axis) and all 8 subjects (colored bars) for variants of the AlexNet and GNet-based encoding models. The error is estimated by sampling with replacement for all estimates of voxelwise validation accuracy and the error displayed is obtained via error propagation.

In contrast, when GNet models were fit independently (“GNet8r-fpf”) there was no decline in the unique or specific contributions of the bottom layers from V1–V4. For V1–V3 the unique contribution of the lower layers was roughly 75%, and for V4 the unique contribution of the lower layers was above 90%. Thus, collectively the GNet8r models do not induce an ordering of tuning with respect to V1–V4, as it is possible to express representations in V1–V4 with the same number of DNN layers (Fig. 6). It is important to recall that the jointly and independently trained GNet models utilized the same architecture and non-linearities, and yield very similar prediction accuracy (Fig. 5c).

### 2.5 Representations in anterior areas do not necessarily entail representations in posterior areas

We first tested for entailment in the AlexNet-based encoding model, because the demonstration of ordered representations in this model strongly suggests that it will also show entailment. We developed a test for entailment based upon transfer learning [Zamir et al., 2018]. First, we trained four independent GNet models (i.e., a GNet feature extractor plus a read-out head) to predict the outputs (i.e., predicted brain activity) of the AlexNet model for V1, V2, V3 and V4, respectively. In other words, we treated the outputs of the AlexNet model as if they were synthetic brain data, and used the GNet8r modeling approach to copy its representations into four distinct networks. The correlation between the outputs of these “reference” GNet models and the outputs of the original AlexNet model was near 1, indicating that we were able to (almost) losslessly copy the AlexNet model representations for each brain area using four independent, stand-alone GNet models. We denote this correlation 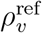, where the subscript indexes voxels. If *v* ∈ *V_i_*, this should be read as the “prediction accuracy of a model with a GNet feature extractor and a read-out head trained and tested on *V_i_*”.

Next, we tested if the representations in each of the reference GNet models were transferable. For each pair of brain areas (*V_i_*, *V_j_*), we froze the feature extractor of the reference model for *V_j_*, and then trained a new read-out head to predict the outputs of the reference model for *V_i_*. We then calculated the prediction accuracy of this new “transfer” model of *V_i_*. We denote this prediction accuracy 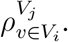. This should be read as the “prediction accuracy of a model with GNet feature extractor trained on *V_j_*, and read-out head trained and tested on a voxel *v* in *V_i_*” (Fig. 7a).

**Figure 7:**
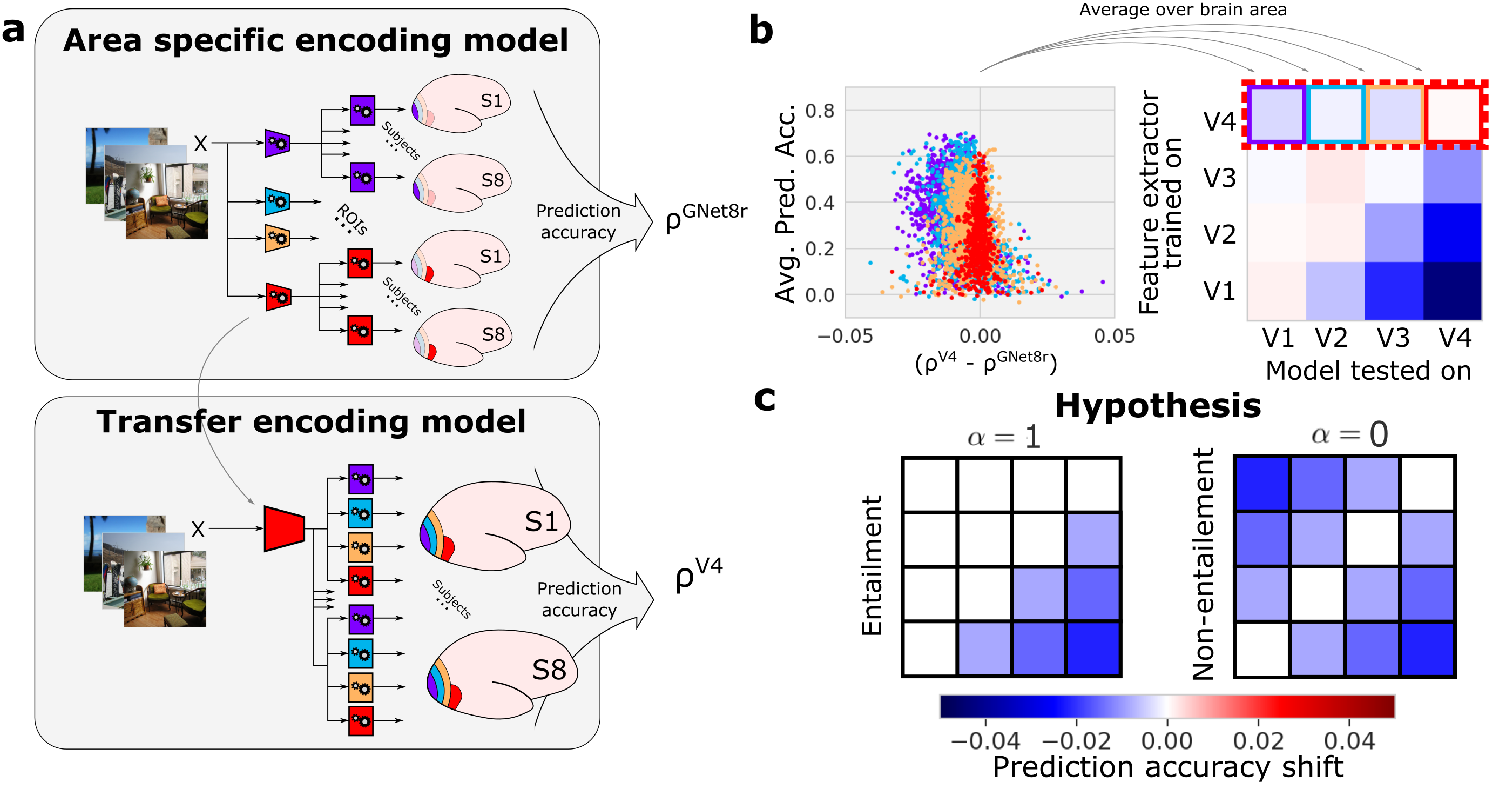
Testing entailment of representations using transfer learning. (**a**) Transfer learning procedure. Stand-alone GNet models consisting of a feature extractor (trapezoids) and a read-out head (squares) are trained (gears) to predict brain activity (shown), or the outputs of another encoding model (not shown) for V1–V4 independently. When trained on brain activity (as shown) the prediction accuracy of these “reference” models is just *ρ*^GNet8r^. The GNet feature extractors of these reference models are used to construct transfer models. For each specific brain area (V4 in this example) the GNet is frozen, and a new read-out head is trained for all areas, yielding prediction accuracy *ρ*^*V*_*j*_^, where the superscript indicates the brain area used to train the feature extractor. In this example *j* = 4 corresponding to area V4. (**b**) To determine how well representations transfer across brain areas, we compute the prediction accuracy shift (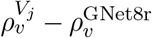, illustrated using an advantage/accuracy plot) for each voxel *v*. For each pair of brain areas *V_j_*, *V_i_* we average these shifts over all voxels *v* ∈ *V_i_*, constructing a prediction accuracy shift matrix (at right is a hypothetical example). Rows of the matrix index the brain area used to train the feature extractor (*V*_j_); columns of the matrix index the brain area over which prediction accuracy shifts are averaged (*V_i_*). (**c**) For representations with entailment (*α* = 1), negative prediction accuracy shifts (blue) accumulate only below the diagonal of the matrix. For representations without entailment (*α* = 0) negative prediction accuracy shifts accumulate above and below the diagonal.

Finally, we constructed a 4 × 4 matrix of differences between the accuracy of reference and transfer models, averaged over all voxels (Fig. 7b). For each element (*i,j*) of the matrix we calculated:

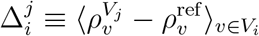

This “prediction accuracy shift matrix” reveals how accurately a DNN that is trained to model the representations in one one brain area can model the representations in another brain area. According to our definition of entailment, the representations in networks trained to model a more posterior area should be a subset of the representations in a network trained to model a more anterior area, but not *vice versa*. Thus, if strict entailment holds, all diagonal and upper-diagonal elements of this matrix should be zero, while all lower-diagonal elements should be negative (Fig. 7c).

As expected, the prediction accuracy shift matrix for the AlexNet model closely resembled the ideal of strict entailment (Fig. 8a). An index we devised to measure resemblance to this ideal had a value of *α* = 1.0 ± 0.2, where *α* = 1 indicates strict entailment, and *α* = 0 indicates no entailment (and *α* = −1 would indicate “reverse” entailment).

**Figure 8:**
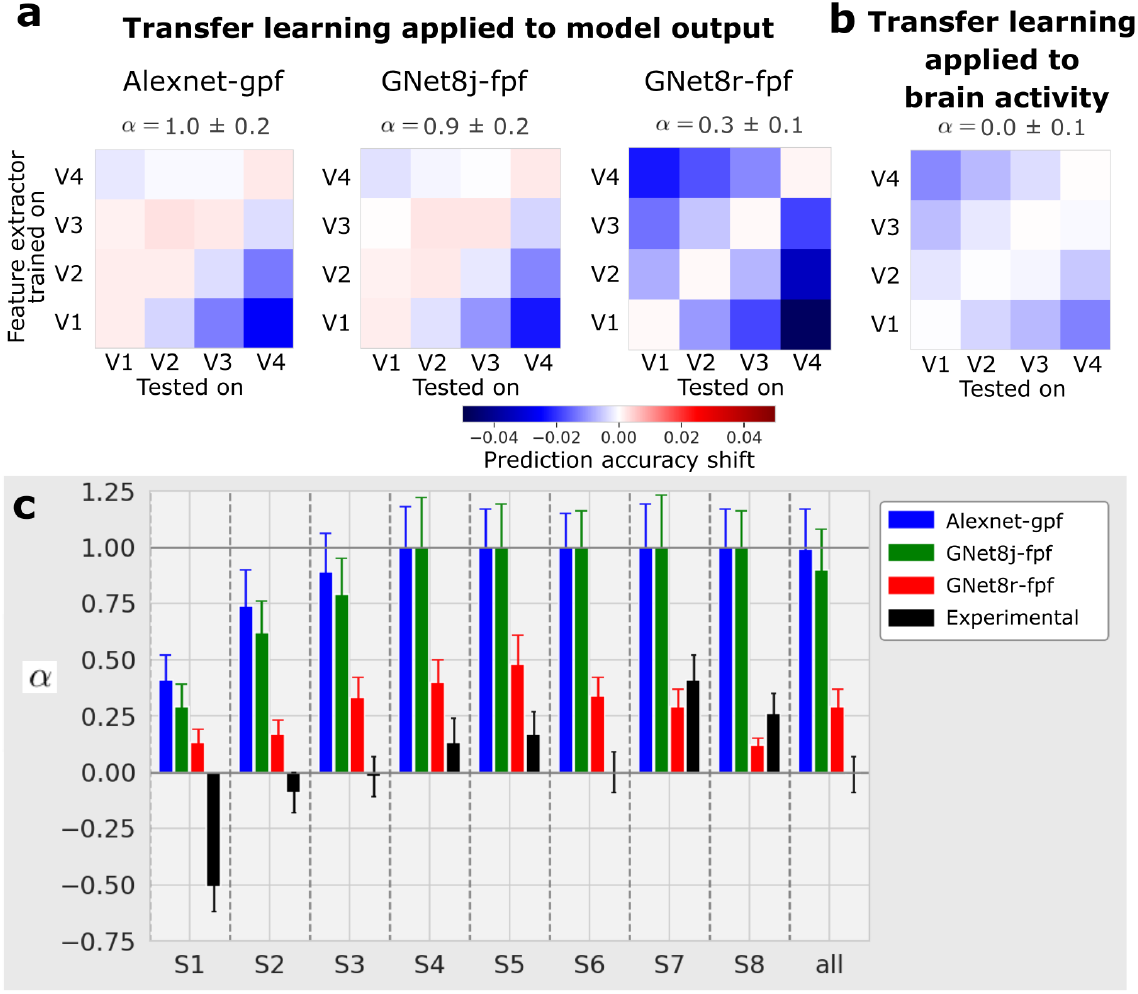
Transfer learning analysis for DNN-based encoding models and V1–V4. (**a**) Transfer learning applied to model outputs. Prediction accuracy shift matrices resulting from the transfer learning procedure applied to outputs of the AlexNet-gpf model, the GNet8j-fpf model, and the GNet8r-fpf model. (**b**) Transfer learning applied to brain activity. Prediction accuracy shift matrix resulting when the transfer learning procedure is applied to measured brain activity. All matrices are averaged across 8 subjects. (**c**) Values of α estimated for individual subjects for the four test cases shown above.

We conducted an identical analysis on the outputs of the jointly trained GNet model because, like the AlexNet-based encoding model, the representations in GNet8j are ordered. Again, and as expected, the prediction accuracy shift matrix closely resembled the ideal for strict entailment, with *α* = 0.9 ± 0.2.

In contrast, when this transfer learning analysis was applied to the outputs of the four GNet models trained independently (GNet8r), the prediction accuracy shift matrix most closely resembled the ideal outcome for non-entailment (*α* = 0.3 ± 0.1). Thus, we established that the GNet8r model learned representations that were neither ordered or entailed with respect to V1–V4.

Finally, we applied the transfer learning analysis to the measured brain activity itself (Fig. 8b and c, black bars). In this case, *ρ*^ref^ = *ρ*^GNet8r^ (as illustrated in Figure 7). To calculate prediction accuracy shifts, we took the GNet optimized to predict V4 brain activity (for instance), froze its parameters, and then re-trained read-out heads to predict brain activity in all areas (Fig. 7). We performed this transfer-learning procedure for feature extractors trained on every brain area. Interestingly, the resulting prediction accuracy shift matrix was most consistent with a non-entailed representation (*α* = 0.0 ± 0.1). It is important to note, however, that when applying transfer learning directly to measured brain activity (as opposed to the outputs of other models), noise can impact the results in potentially complicated ways that make interpretation difficult.

## 3 Discussion

We demonstrated that brain-optimized networks can be made to yield predictions of brain activity that are more accurate for most voxels than predictions read-out from task-optimized networks. We then used brain-optimized networks to test for evidence that hierarchical representations are necessary to achieve this level of prediction accuracy, focusing specifically on the intuitive properties of *ordering* and *entailment*. First, we showed that four GNets trained independently on V1–V4 (GNet8r) predicted brain activity as accurately as a single GNet trained jointly on V1–V4 (GNet8j), thus demonstrating that a training strategy biased in favor of hierarchical representations offered no advantage in prediction accuracy over a training strategy that was equally amenable to non-hierarchical representations (GNet8r). We then showed that in encoding models based on AlexNet and GNet8j, lower layers contributed most strongly to posterior areas (ordering), and representations optimized for anterior areas were more readily transferable to posterior areas than the other way around (entailment). In contrast, for GNet8r lower layers contributed equally to posterior and anterior areas, and optimized representations showed the same prediction accuracy shift when transferred to anterior or posterior areas.

These results have important implications for the ongoing debate about how literally DNNs can be interpreted as mechanistic models of the primate visual system [Kriegeskorte, 2015, Kay,2018, Richards et al., 2019, Lindsay, 2021, Cao and Yamins, 2021]. Clearly, the connections between layers in the kind of feed-forward DNNs we have studied here cannot be interpreted as literal stand-ins for anatomical connections in the brain; the best such networks can do is model stimulus-dependent representations encoded in brain activity patterns that are structured by far more complicated recurrent circuits. However, our work has allowed us to interrogate a key assumption in many models and theories of primate vision: that visual brain areas, like layers in a DNN, are related through hierarchical composition of a fixed computation.

Taken literally, the assumption of hierarchical composition implies that each visual brain area computes a fixed, or “canonical” computation on their inputs. As in a DNN, the outputs of this computation are sent to the brain area above and the inputs to it are received from the brain area below. Although the canonical computation may not be identical across areas, it has a fixed form: across areas the computation performed varies only up to a set of parameters values θ. The assumption of hierarchical composition also implies that the canonical computation in the brain is “simple”, in the sense that variation in representations across V1–V4 cannot be explained by variation in *θ* in one layer alone. Instead, variation in representations across V1–V4 can only be explained by varying compositional depth.

We did not directly test these implications, but we note that it is currently unclear if any of them are true. Efforts to articulate a canonical computation in cortical circuits has been ongoing since the early descriptions of cortical organization [Mountcastle, 1998, Hubel and Wiesel,1977], and it is still not known if this concept will prove to be essential for understanding cortical function. It is also not clear if any putative canonical computation in the brain will turn out to be “simple” in the sense defined above. Recent studies suggest that the input/output mapping of single cortical neurons may be quite complex [Beniaguev et al., 2021, Poirazi et al.,2003, Gidon et al., 2020].

We were, however, able to directly test two additional and no-less critical implications of hierarchical composition: that accurate models of V1–V4 should exhibit ordering and entailment. Ordering arises directly from the assumption that variance in brain representations across areas can only be explained by varying compositional depth. Entailment arises as a consequence of composition and implies that one cannot build a DNN model of V4 without also building a DNN model of V1–V3, and so on.

Our strategy for testing these implications of hierarchical composition was to demonstrate the existence of a DNN that predicts as accurately as any other DNN-based encoding model, using the same nonlinearities and architecture, but does not show ordering or entailment. We used brain-optimization to construct such a counterexample, GNet8r. The fact that GNet8r exhibits neither ordering nor entailment indicates that these properties are not essential for obtaining accurate predictions of brain activity. It follows that the ordering of V1–V4 with respect to layers in many task-optimized networks is an outcome that is contingent on specific model-building choices (e.g., selection of training samples, task-optimization vs. brain-optimization, and network constraints on width and expressivity). It further follows that the depth of the network layer to which a brain area most closely aligns is not a reliable proxy for the “complexity” of representations in these brain areas. At best, the network layer most closely aligned with a single brain area is an upper bound on the complexity of representations in the brain area, and our work shows that the upper bound estimated from any single DNN-based encoding model can be quite loose. Our results thus encourage caution in the interpretation of DNNs as mechanistic models of the visual system [Kay, 2018]. Although DNNs will continue to be useful tools for inspiring and exploring brain models, it is currently unclear what aspects of DNNs are specifically brain-like.

Why do networks that do not show ordering and entailment yield the same prediction accuracy as those that do? One possible explanation may have to do with the difficulty of accounting for all of the explainable variance in brain activity. The network-based models we constructed do not account for between 35 to 55% of the explainable variance in V1–V4 (Fig. 3). Thus, networks that vary with respect to ordering and entailment may simply account for different portions of the variance in brain activity. Another possibility is that DNNs are, as a family of models, simply degenerate with respect to many of the properties that visual neuroscientists currently consider interesting and important. Indeed, it is increasingly appreciated that DNNs with widely varying architectures [Storrs et al., 2021, Xu and Vaziri-Pashkam, 2021] afford similar levels of performance in predicting brain activity. Relatedly, even though early results suggested that supervised training was required for accurate correspondence between networks and brain activity [Khaligh-Razavi and Kriegeskorte, 2014], recent advances have shown that unsupervised models can explain brain activity just as well [Zhuang et al., 2021].

Finally, our results have an interesting implication for how to understand the function of V1–V4. In a strict hierarchical interpretation of V1–V4, each area functions effectively as a pre-processing unit that participates in the sequential construction of a representation expressed in some more anterior area. Under this interpretation, if the more anterior brain areas fed by V1–V4 were damaged, V1–V4 would no longer have any functional role. Furthermore, if representations in an earlier area were changed (e.g., via a perceptual learning task), representations in all anterior areas would presumably have to change as well. Distributing representational labor across the layers of a strict hierarchy is understood to be the essential for the successes of deep neural networks trained with stochastic gradient descent, as in many cases, it permits representing complex relations with less neural units [Bahri et al., 2020]. However, this form of labor distribution strikes us as a precarious and inefficient arrangement for biological networks. In contrast, if V1–V4 was not a strict hierarchy, each area could in principle encode representations that are optimized for dedicated, independent tasks, or could encode representations that are routinely combined in non-hierarchical ways to solve novel tasks as they emerge.

We interpret this work as a corrective to a tendency to treat “hierarchy” as the defining feature of primate visual organization. Our interpretation is consistent with emerging evidence for non-hierarchical representation in other sensory systems [Hamilton et al., 2021], motivates the development of networks with multi-branch architectures [Bakhtiari et al., 2021] that model parallel visual streams with distinct functions [Ungerleider, 1982, Schiller and Logothetis, 1990,Pitcher and Ungerleider, 2021], and underscores the need for computational models that treat hierarchy as an emergent property [Konkle, 2021] rather than a requirement for successful vision.

## 4 Methods

### 4.1 Dataset acquisition in brief

All models were trained on the Natural Scenes Dataset (NSD). Complete details on NSD are provided elsewhere [Allen et al., 2022]. Briefly, the NSD dataset consists of between 22K and 30K fMRI image-responses per subject (8 subjects). Images were sampled from the Common Objects in Context (COCO) database [Lin et al., 2015] and displayed at 8.4° × 8.4°. The experimental design specified that each of the eight participants would view 10K distinct images (3 presentations each), and a special subset of 1K images would be shared across participants (8 subjects × 9K unique images + 1K shared images = 73K unique images); however, not all participants completed the full acquisition, so the final numbers are somewhat smaller. All fMRI data in the NSD were collected at ultra-high field (7T) using a whole-brain, 1.8-mm, 1.6-s, gradient-echo, echo-planar imaging (EPI) pulse sequence.

The image-responses are expressed in terms of betas obtained from a general linear model (GLM) analysis. For this paper, we used GLM results provided with the NSD data release, specifically, the 1.8-mm volume preparation of the data and version 3 of the GLM betas (be-tas_fithrf_GLMdenoise_RR). This GLM version involves estimating the hemodynamic response function for each voxel, using the GLMdenoise technique for denoising [Kay et al., 2013a], and using ridge regression to improve the estimation of single-trial betas. Betas indicate BOLD response amplitudes evoked by each stimulus trial relative to the baseline signal level present during the absence of a stimulus (“gray screen”). The betas for each voxel in each session were separately z-scored and all sessions were concatenated.

### 4.2 NSD synthetic experiment

In addition to the core NSD experiment, the 8 subjects also participated in an additional 7T scanning session termed ‘nsdsynthetic’. This session involved presentation of a variety of artificial stimuli. Procedures for data acquisition, pre-processing, and GLM analysis were the same as for the NSD core. Stimuli consisted of a set of 284 images that can be conceptually grouped as follows (the number of distinct images in each group is indicated in parentheses): white noise (4), white noise with a large block size (4), pink noise (4), natural scenes (4), upside-down versions of these scenes (4), Mooney versions of these scenes (4), line-drawing versions of these scenes (4), contrast-modulated natural scenes (4 scenes × 5 contrast levels (100%, 50%, 10%, 6%, 4%) = 20), phase-coherence-modulated natural scenes (4 scenes × 4 coherence levels (75%, 50%, 25%, 0%) = 16), single words (4 words × 5 positions × 2 word lengths = 40), spiral gratings varying in orientation and spatial frequency (112), and chromatic pink noise varying in hue (68). Images typically occupied 8.4°× 8.4° (same as NSD core), though a few of the word stimuli extended beyond this extent. Examples of the stimuli are provided in Supplementary Figure 1.

Stimuli were presented in pseudorandom order using a 2-s ON/2-s OFF trial structure. Stimuli were shown against a gray background with an RGB value of (126, 110, 108), and were delivered using a linear color lookup table. During each run, a small semi-transparent gray fixation dot with a black border (0.2° × 0.2°, 50% opacity) was present at the center of the stimuli. The luminance of the dot changed every 1.4 s. In alternating runs, while maintaining central fixation, subjects either performed a fixation task (report direction of the luminance change of the fixation dot) or a one-back task (report whether the current image is the same as the previous image). A total of 8 runs (each with duration 428 s) were collected, yielding a total of 744 stimulus trials over the course of the scan session. For the analyses performed in this paper, we modeled each stimulus trial (ignoring the variations in task performed by the subject) and considered only the central 8.4°square region (matching NSD core).

### 4.3 Identification of visual brain areas V1–V4

Human visual brain areas were identified using a separate population receptive field (pRF) retinotopic mapping experiment, as documented in [Allen et al., 2022]. Retinotopic areas (more generally described as ‘regions of interest’ (ROIs)) were manually drawn based on results of the pRF experiment. These ROIs consist of V1v, V1d, V2v, V2d, V3v, V3d, and V4, and extend from the fovea (0° eccentricity) to peripheral locations that exhibit sensible responses in the pRF experiment given the limited stimulus size (the diameter of the pRF mapping stimulus was 8.4°). The total number of voxels (cumulative over subjects) in each ROI was 9041, 8818, 7763 and 3975 for V1, V2, V3 and V4 respectively, totaling 29597 voxels, with one subject contributing as few as 3027 voxels, and another as many as 4627 voxels.

### 4.4 General encoding model architecture

Encoding models based upon both the task-optimized and brain-optimized networks consisted of a feature extractor (a DNN) and multiple read-out heads as detailed in the following sections and described in Allen et al. [2022].

#### 4.4.1 Feature extractor

Feature extractors for all encoding models are sequences of transformations

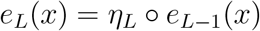

operating on *x*, here an input image, where *η_L_* is the transformation that operates at layer *L* on the output of the subsequence *e*_*L*−1_(*x*). *e*_*L*−1_(*x*) and *η_L_* may themselves denote arbitrary sequences of transformations. Our encoding models leverage the multiple intermediate representations *e_l_*(*x*), which are feature maps whose elements are denoted by [*e_l_*(*x*)]_*kji*_, where *k* indexes features and (*i, j*) are pixel coordinates in each feature map.

#### 4.4.2 Read-out heads

Read-out heads convert features output by the feature extractor into predictions of brain activity, 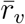, for each voxel *v*. These predictions can be expressed as a linearized model

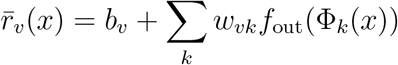

where

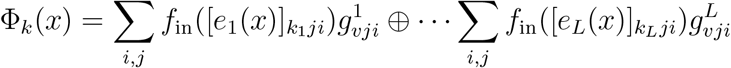

and where *f_in_*(·) and *f_out_*(·) are typically some compressive nonlinearity and the sum ⊔ denotes the concatenation along the feature axis *k* = (*k*_1_,… *k_L_*). 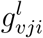 is the value of the “pooling field” for voxel *v* applied at feature maps in layer *l* at pixel (*i,j*). Pooling fields generalize the population receptive field [Dumoulin and Wandell, 2008] to arbitrary feature maps. All pooling field elements are positive-valued and normalized such that their sum equals to unity. Figure 2 illustrates the two pooling field variants used in our work. Gaussian pooling fields (gpf) are fully described by three parameters that specify the position and size of a symmetric 2D Gaussian function. For “flexible” pooling fields (fpf) each pixel value of the pooling field is an independent and learnable parameter.

### 4.5 AlexNet encoding model

#### 4.5.1 Feature extractor

For the task-optimized model featured in all figures of the main text, the feature extractor was an AlexNet deep convolutional neural network trained to classify 1000 object categories of the ImageNet database Krizhevsky et al. [2012]. We used the pre-trained weights from Torchvision’s model zoo (https://pytorch.org/vision/stable/models.html). Not all feature maps were used in the encoding. At each layer specified in Supplementary Table 3, if a layer had more than 512 feature maps, we selected the 512 feature maps with the most variance with respect to the COCO images in our experiment. The final model thus exposed a total of 2688 feature maps.

#### 4.5.2 Read-out head

We constructed two variants of the AlexNet model: one with a Gaussian pooling field (AlexNet-gpf), and one with a flexible pooling field (AlexNet-fpf). To construct AlexNet-fpf models, feature maps with the same spatial resolution were concatenated and a distinct spatial pooling field was learned for each spatial resolution. Thus, for AlexNet-fpf models the read-out heads included multiple pooling fields. For both model variants, feature maps from each layer throughout the depth of the AlexNet feature extractor were input to the read-out head. For details see Supplementary Table 3.

#### 4.5.3 Training

For AlexNet models the parameters of the feature extractor were pre-trained, as described above. Thus, only the parameters of the read-out heads were optimized.

For AlexNet-gpf models the three pooling field parameters are learned via grid search over a list of 2680 candidates tiling the visual field with 8 log-spaced sizes varying from 3 to 40% of stimulus size and spaced roughly in proportion to their sizes (such that each size tiles the visual field fully). For each candidate receptive field, the tuning weights are learned via ridge regression with the ridge parameter selected to maximize validation accuracy on a held-out 10% set of the training set.

For AlexNet-fpf models the training of the read-out heads was performed via gradient descent with the ADAM optimizer (lr = 10^−3^, *β*_1_ = 0.9, *β*_2_ = 0.999).

### 4.6 GNet encoding model

#### 4.6.1 Feature extractor

We refer to the feature extractor of the brain-optimized network as a “GNet”. The GNet feature extractor consists of a pre-filtering network (*e*_1_(*x*)) followed by a deep feature extractor.

The input image resolution of the pre-filtering network is 227 × 227 and outputs an embedding of 192 features at a 27 × 27 spatial resolution. This part of the network is identical to the first two layers AlexNet.

The deep feature extractor is a sequence of blocks of layers. Each block consists of a batch normalization layer followed by a dropout layer and a convolutional layer that preserves feature map resolution. The use of batch normalization followed by dropout in each block has been characterized as performing input whitening and decorrelation Chen et al. [2019]. After three blocks, the resolution of the feature maps is reduced from 25 × 25 to 13 × 13 by a max pooling layer. The GNet sequence of layers was designed so that each pixel in the final convolutional layer effectively pools over the whole input image, whereas pixels in lower layers only integrate over more local regions of the input image.

#### 4.6.2 Read-out head

For the GNet model only flexible pooling fields were used due to training requirement. As with the AlexNet model, feature maps with the same spatial resolution were concatenated and a single spatial pooling field was learned for each spatial resolution. Thus, the read-out heads contained multiple pooling fields.

Read-out heads employed a fully differentiable nonlinearity:

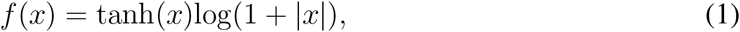

that was applied either before or after spatial pooling, or both. This nonlinearity has several interesting and desirable characteristics: 1) it has an expansive and a compressive regime, 2) it is differentiable everywhere, with no discontinuity and 3) it does not plateau over a large range. The final GNet model benefited from using this nonlinearity before and after spatial pooling (i.e. as *f_in_* and *f_out_*).

During the structure and hyperparameter selection process for GNet, we noticed that the best extant GNet model was obtained when some feature maps in lower layers were not directly connected to the read-out heads, as indicated on Supplementary Table 3. Note, however, we find that the results of analyses reported here are unchanged when using fully connected read-out heads.

#### 4.6.3 Training

The pre-filtering network was taken from a task-optimized AlexNet. Its parameters were kept fixed during brain-optimization. While it was possible to optimize the pre-filtering network parameters along with the deep extractor parameters from a random initial condition, the results using a pre-trained filtering network were slightly better for all model variants.

All parameters of the deep feature extractor and the read-out heads are learned jointly via gradient descent with the ADAM optimizer (lr = 10^−3^, *β*_1_ = 0.9, *β*_2_ = 0.999). The training steps of the deep feature extractor and the read-out heads are alternated to promote stability of the training procedure.

To account for the noise profile and the discrepancies in voxel predictability, we used the following *L*2-norm weighted loss function

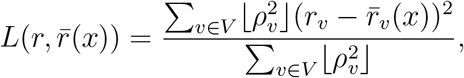

where 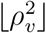 is the batchwise Pearson-like correlation of every voxel with a floor of 0.1 to always permit some contribution of the yet-to-be-predicted voxels (i.e. those with low correlation at the onset of training) in the voxel ensemble *V*. If the model can be trained voxelwise, then this weighting could be ignored but this could affect the learning dynamics (learning rates).

Unlike a typical deep network optimization with gradient descent, where all parameters subserve the global objective, this problem involves shared parameters (feature extractor) and voxelwise parameters (read-out heads). A global early stopping criterion may not be optimal for all voxels, since overfitting would be detected when the average loss starts to increase. On the other hand, voxelwise loss may increase before the optimal point due to the dependence on the changing deep feature extractor. This remains an outstanding problem. In spite of the aforementioned issues, relative successes were achieved by using a global early stopping criterion with the parameters clamped to their best value according to the validation accuracy of a 10% holdout set of the training set.

A second challenge emerged from the fact that each read-out head only has access to the brain responses associated with the subject-wise image samples, while the feature extractor leverages all available images. We addressed this issue by interleaving samples from each subject. While typical training consists of randomly sampling a minibatch from the training set, we first selected a subject at random and sampled a minibatch from its set of training images and brain responses. The read-out heads follow the matching brain response voxels and changes from batch to batch while the shared feature extractor is trained with gradients backpropagated from the current read-out heads. At the end of each epoch, every sample from every subject has been “seen” exactly once. This procedure yielded distinct improvements in model accuracy relative to subject-wise training, i.e. training a feature extractor network for each subject separately (as in Allen et al. [2022]).

In some cases, we also performed a second and third phase of training (which we refer to as “fine tuning”, GNet8jft-fpf in Fig. 3e). Each subsequent phase restarts training from the optimal weight values from the previous phase. The second phase follows the same procedure as the first phase but all the read-out heads parameters were fixed and only the feature extractor parameters were trained (including those of the pre-filtering network). This second phase accounted for most of the improvement over the first phase. A third phase followed again the same objective in which we trained only the parameters of the read-out heads, while the feature extractor remained constant.

### 4.7 Gabor encoding model

#### 4.7.1 Feature extractor

The Gabor model feature extractor consists of a single fixed set of convolutions: 12 Gabor wavelets with spatial frequency log-spaced between 3 and 72 cyc/stimulus at 6 evenly-spaced orientations between 0 and *π*.

#### 4.7.2 Read-out head

We used a read-out head with a Gaussian pooling field for the Gabor model. Following previous work [St-Yves and Naselaris, 2018], we used a compressive nonlinearity *f_in_*(*x*) = log(1 + |*x*|) while *f_out_*(*x*) = *x*.

#### 4.7.3 Training

Gabor models were fit using grid search over the pooling field parameters (same candidate grid as for the AlexNet-gpf model), followed by ridge regression to determine the feature weights.

### 4.8 Cross-validated prediction accuracy

Prediction accuracy is calculated using the subset of trials that include images displayed to all subjects (for most subjects, 1000 images with 3 repeats each). These trials were not included in the training data, and were not used for hyperparameter selection (see below), ensuring proper cross-validation. Prediction accuracy is the Pearson correlation between predicted brain activity and measured brain activity on a single-trial basis. Prediction accuracy uncertainty was estimated by sampling with replacement the predicted and actual brain activities for each voxel.

### 4.9 Hyperparameter selection

The process of hyperparameter selection for the GNet models (e.g., pooling map resolutions, number of layers, nonlinearities, forms of regularization) was based on the prediction accuracy measured on a fixed, held-out model selection set consisting of 10% of each subject’s training data. The holdout prediction accuracy was evaluated, for each selection of hyperparameter values, at the early stopping point (i.e. at the minimum of the holdout loss during training of that model). The GNet model was refined on this basis using 4 subjects, and the best extant model was used to perform the final analysis with all 8 subjects.

### 4.10 Receptive field modeling

Receptive fields (Fig. 4) were derived directly from the spatial pooling fields of the read-out head for each voxel. For some voxels, the flexible pooling fields do not have a clearly localized structure. However, for the vast majority of voxels the trained flexible spatial pooling fields extend smoothly from a clearly identifiable center. Thus, we characterized these maps by fitting an elliptical 2D gaussian to the parameters (the pixels in the map) of the flexible spatial pooling field. For each voxel, the size of its receptive field is defined as 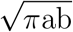, where *a* and *b* are one std. dev. along the major and minor axis of the elliptical gaussian, and its eccentricity is the Euclidean distance from the display center. In the plot of Figure 4d both measures are expressed as percent of the display size (i.e. 100% ≡ 8.4°).

### 4.11 Layer-wise contributions to prediction accuracy

To test for alignment between layer depth and brain areas we calculated the prediction accuracy of network-based encoding models when layers from the bottom or top half of the feature extractor network were masked. Specifically, let *ρ*_bottom_ be the prediction accuracy obtained from an encoding model in which feature weights *w* from layers in the top half of the feature extractor network have been zeroed-out (refer to Figure 6a for the designation of bottom and top layers in the AlexNet and GNet feature extractors, respectively). Separately, we also calculate the prediction accuracy *ρ*_top_ obtained by zeroing-out the feature weights that couple to the bottom layers. Both of these measures of prediction accuracy are necessarily smaller than the *total* prediction accuracy of the model ρ obtained when all feature weights are used (see Venn diagram in Figure 6b).

To assess ordering for a given DNN, we calculated the specific and unique contributions of the bottom layers to predicting brain activity. Given that total variance varies a great deal across voxels, we express the specific and unique contributions of the bottom layers for each voxel in relative terms:

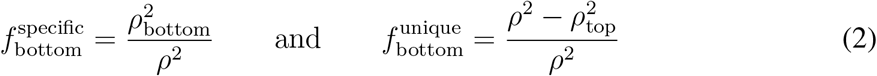

This decomposition of prediction accuracy into specific and unique contributions is performed for individual voxels. We then average the specific and unique contributions over all voxels in a single brain area that have validation accuracy that exceeded *ρ* = 0.055 (*p* < 0.01, prediction randomization trial). We plot these values along the presumed hierarchy of brain areas in Figure 6c. We verified that the results were robust to the precise choice of threshold. Note that values for the specific and unique contribution of the top layer are obtained by swapping “bottom” and “top” subscripts in the above formulas, and can be read-off by eye from the curves in Figure 6c.

### 4.12 Transfer learning experiments

In transfer learning experiments, we test how well representations in a GNet feature extractor optimized for one brain area can generalize to another brain area. To test this, we first train a GNet feature extractor and read-out head to predict brain activity (or the outputs of another encoding model) for all voxels in a single brain area i. We then freeze the parameters of the trained GNet feature extractor for this area-specific encoding model (we call it the “reference model” below), but train new read-out heads to predict brain activity in all brain areas. We then compare the cross-validated prediction accuracy obtained when the feature extractor and read-out are trained on the same brain area (the reference model) to prediction accuracy obtained when the feature extractor and read-out are trained on different brain areas (the “transfer” model).

Let *i,j* ∈ (1, 2, 3, 4) index brain areas, and let 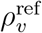 be the cross-validated prediction accuracy of a reference model for voxel *v*. For the reference model, the GNet feature extractor and the read-out head are trained simultaneously on the brain area that *v* belongs to and then used to predict activity on held-out trials for that area. Let 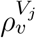 be the cross-validated prediction accuracy of a transfer model. For the transfer model, the feature extractor is trained on area *V*j** and then frozen while a read-out head is subsequently trained on the brain area to which *v* belongs. The transfer model is then used to predict activity on held-out trials in the brain area to which *v* belongs. We define the “prediction accuracy shift” 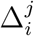 as:

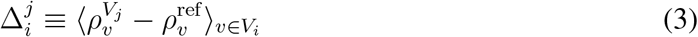

where 〈·〉_*v*_ ∈*V_i_* denotes averaging over all voxels in brain area *V_i_*. In Figures 7 and 8 we show prediction accuracy shift matrices consisting of 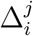 for all pairs of brain areas *V_i_*, *V_j_*. Rows (labeled “feature extractor trained on”) and columns (labeled “model tested on”) correspond to the superscript and subscript of 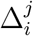, respectively. For entries along the diagonal of this matrix, 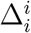 measures the prediction accuracy shift induced by first training the feature extractor and read-out head simultaneously on area *V_i_* (i.e., the reference model) and then fixing the feature extractor and re-training the read-out head only on area *i* (i.e., the transfer model). The subtle difference between simultaneous and serial training of the model components typically induces a small positive prediction accuracy shift.

In Figure 7 we illustrate the transfer of a GNet feature extractor optimized for V4 to V1,V2 and V3. The results of this transfer learning operation populate the top row, 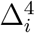, of the prediction accuracy shift matrix.

In Figure 8a, the three prediction accuracy shift matrices on the left labeled “AlexNet-gpf”, “GNet8j-fpf” and “GNet8r-fpf” were constructed by applying the transfer learning procedure just described to the outputs of the AlexNet-gpf, GNet8j-fpf and GNet8r-fpf models, respectively. In other words, instead of using brain activity to train the reference and transfer models, we use the outputs of the indicated encoding models to train them (by “outputs”, we specifically mean the activity predictions of the encoding model for each voxel). This allows us to identify the models that learn hierarchical representations during the course of their training. For the matrix on the far right labeled “Measured brain activity”, the transfer learning procedure was applied directly to measured brain activity. Note that in this special case, 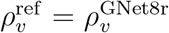 for all *v*.

We characterized the structure of the prediction accuracy shift matrices by calculating a scalar value, *α*, that captures the normalized difference between the upper and lower triangular components:

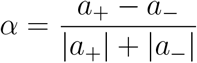

where *a*_+_ and *a*___ are the slopes of the upper and lower triangular entries of the matrix, respectively. To calculate these slopes, we expressed the respective matrix entries as a function of their taxicab distance from the matrix diagonal, the off-diagonal entries having a distance of 1 and so on. In cases where the matrix is characterized by a salient lower triangular (negative slope) and a flat (zero slope) upper triangular, *α* would be close to 1. On the other hand, if the matrix is symmetric, *α* would be zero. The error in *α* is estimated via error propagation of the estimate of the errors on the slopes.

## 5 Acknowledgements

This work was supported by NSF CRCNS grants IIS-1822683 (K.K.) and IIS-1822929 (T.N.).

## 6 Author contributions

G.S.-Y. and T.N. conceived of the project and wrote the paper. E.J.A. and Y.W. performed data collection for the Natural Scenes Dataset. G.S.-Y. performed data analysis. G.S.-Y., T.N. and K.K. discussed and interpreted results. T.N. directed the overall project. All authors discussed and edited the paper.

## References

E. J. Allen, G. St-Yves, Y. Wu, J. L. Breedlove, J. S. Prince, L. T. Dowdle, M. Nau, B. Caron, F. Pestilli, I. Charest, et al. A massive 7t fmri dataset to bridge cognitive neuroscience and artificial intelligence. Nature neuroscience, 25(1):116–126, 2022.

J. Antolík, S. B. Hofer, J. A. Bednar, and T. D. Mrsic-Flogel. Model constrained by visual hierarchy improves prediction of neural responses to natural scenes. PLoS computational biology, 12(6):e1004927, 2016.

Y. Bahri, J. Kadmon, J. Pennington, S. S. Schoenholz, J. Sohl-Dickstein, and S. Ganguli. Statistical mechanics of deep learning. Annual Review of Condensed Matter Physics, 11(1): 501–528, 2020. doi: 10.1146/annurev-conmatphys-031119-050745.

S. Bakhtiari, P. Mineault, T. Lillicrap, C. Pack, and B. Richards. The functional specialization of visual cortex emerges from training parallel pathways with self-supervised predictive learning. 2021.

E. Batty, J. Merel, N. Brackbill, A. Heitman, A. Sher, A. Litke, E. Chichilnisky, and L. Paninski. Multilayer recurrent network models of primate retinal ganglion cell responses. 2016.

D. Beniaguev, I. Segev, and M. London. Single cortical neurons as deep artificial neural networks. Neuron, 109(17):2727–2739, 2021.

S. A. Cadena, G. H. Denfield, E. Y. Walker, L. A. Gatys, A. S. Tolias, M. Bethge, and A. S. Ecker. Deep convolutional models improve predictions of macaque v1 responses to natural images. PLoS computational biology, 15(4):e1006897, 2019.

R. Cao and D. Yamins. Explanatory models in neuroscience: Part 1–taking mechanistic abstraction seriously. arXiv preprint arXiv:2104.01490, 2021.

M. Carandini, J. B. Demb, V. Mante, D. J. Tolhurst, Y. Dan, B. A. Olshausen, J. L. Gallant, and N. C. Rust. Do we know what the early visual system does? Journal of Neuroscience, 25 (46):10577–10597, 2005. ISSN 0270-6474.

G. Chen, P. Chen, Y. Shi, C. Hsieh, B. Liao, and S. Zhang. Rethinking the usage of batch normalization and dropout in the training of deep neural networks. CoRR, abs/1905.05928, 2019.

R. M. Cichy, A. Khosla, D. Pantazis, A. Torralba, and A. Oliva. Comparison of deep neural networks to spatio-temporal cortical dynamics of human visual object recognition reveals hierarchical correspondence. Scientific reports, 6(1):1–13, 2016.

J. J. DiCarlo and D. D. Cox. Untangling invariant object recognition. Trends in cognitive sciences, 11(8):333–341, 2007.

S. Dumoulin and B. Wandell. Population receptive field estimates in human visual cortex. NeuroImage, 39(2):647–660, 2008.

M. Eickenberg, A. Gramfort, G. Varoquaux, and B. Thirion. Seeing it all: Convolutional network layers map the function of the human visual system. NeuroImage, 152:184–194, 2017.

D. J. Felleman and D. C. Van Essen. Distributed hierarchical processing in the primate cerebral cortex. Cerebral cortex (New York, NY: 1991), 1(1):1–47, 1991.

K. Fukushima. Neocognitron: A hierarchical neural network capable of visual pattern recognition. Neural networks, 1(2):119–130, 1988.

A. Gidon, T. A. Zolnik, P. Fidzinski, F. Bolduan, A. Papoutsi, P. Poirazi, M. Holtkamp, I. Vida, and M. E. Larkum. Dendritic action potentials and computation in human layer 2/3 cortical neurons. Science, 367(6473):83–87, 2020.

A. Goyal and Y. Bengio. Inductive biases for deep learning of higher-level cognition. arXiv preprint arXiv:2011.15091, 2020.

K. Grill-Spector and R. Malach. The human visual cortex. Annual Review of Neuroscience, 27 (1):649–677, 2004.

U. Güçlü and M. A. J. van Gerven. Deep Neural Networks Reveal a Gradient in the Complexity of Neural Representations across the Ventral Stream. The Journal of neuroscience : the official journal of the Society for Neuroscience, 35(27):10005–14, jul 2015. ISSN 1529-2401.

L. S. Hamilton, Y. Oganian, J. Hall, and E. F. Chang. Parallel and distributed encoding of speech across human auditory cortex. Cell, 184(18):4626–4639, 2021.

K. A. Hansen, K. N. Kay, and J. L. Gallant. Topographic Organization in and near Human Visual Area V4. Journal of Neuroscience, 27(44):11896–11911, 2007. ISSN 0270-6474.

D. Hassabis, D. Kumaran, C. Summerfield, and M. Botvinick. Neuroscience-inspired artificial intelligence. Neuron, 95(2):245–258, 2017.

L. Henriksson, L. Nurminen, A. Hyvarinen, and S. Vanni. Spatial frequency tuning in human retinotopic visual areas. Journal of Vision, 8(10):5–5, 2008. ISSN 1534-7362.

C. C. Hilgetag and A. Goulas. ‘hierarchy’in the organization of brain networks. Philosophical Transactions of the Royal Society B, 375(1796):20190319, 2020.

K. D. Himberger, H.-Y. Chien, and C. J. Honey. Principles of temporal processing across the cortical hierarchy. Neuroscience, 389:161–174, 2018.

D. H. Hubel and T. N. Wiesel. Receptive fields, binocular interaction and functional architecture in the cat’s visual cortex. The Journal of physiology, 160(1):106–154, 1962.

D. H. Hubel and T. N. Wiesel. Ferrier lecture-functional architecture of macaque monkey visual cortex. Proceedings of the Royal Society of London. Series B. Biological Sciences, 198(1130): 1–59, 1977.

K. N. Kay. Principles for models of neural information processing. NeuroImage, 180:101–109, 2018. ISSN 1053-8119. New advances in encoding and decoding of brain signals.

K. N. Kay, A. Rokem, J. Winawer, R. F. Dougherty, and B. a. Wandell. GLMdenoise: a fast, automated technique for denoising task-based fMRI data. Frontiers in neuroscience, 7(December):247, jan 2013a. ISSN 1662-4548.

K. N. Kay, J. Winawer, A. Mezer, and B. a. Wandell. Compressive spatial summation in human visual cortex. Journal of neurophysiology, 110(2):481–94, jul 2013b. ISSN 1522-1598.

S.-M. Khaligh-Razavi and N. Kriegeskorte. Deep supervised, but not unsupervised, models may explain it cortical representation. PLoS computational biology, 10(11):e1003915, 2014.

W. F. Kindel, E. D. Christensen, and J. Zylberberg. Using deep learning to probe the neural code for images in primary visual cortex. Journal of vision, 19(4):29–29, 2019.

D. A. Klindt, A. S. Ecker, T. Euler, and M. Bethge. Neural system identification for large populations separating what and where. In Proceedings of the 31st International Conference on Neural Information Processing Systems, pages 3509–3519, 2017.

E. Kobatake and K. Tanaka. Neuronal selectivities to complex object features in the ventral visual pathway of the macaque cerebral cortex. Journal of neurophysiology, 71(3):856–867, 1994.

T. Konkle. Emergent organization of multiple visuotopic maps without a feature hierarchy. bioRxiv, 2021.

N. Kriegeskorte. Deep Neural Networks: A New Framework for Modeling Biological Vision and Brain Information Processing. Annual review of vision science, 1(1):417–446, nov 2015. ISSN 2374-4650. doi: 10.1146/annurev-vision-082114-035447.

A. Krizhevsky, I. Sutskever, and G. E. Hinton. Imagenet classification with deep convolutional neural networks. Advances in neural information processing systems, 25:1097–1105, 2012.

Y. LeCun, B. Boser, J. S. Denker, D. Henderson, R. E. Howard, W. Hubbard, and L. D. Jackel. Backpropagation applied to handwritten zip code recognition. Neural computation, 1(4): 541–551, 1989.

Y. Lecun, Y. Bengio, and G. Hinton. Deep learning. Nature, 521(7553):436–444, 5 2015. ISSN 0028-0836. doi: 10.1038/nature14539.

T.-Y. Lin, M. Maire, S. Belongie, L. Bourdev, R. Girshick, J. Hays, P. Perona, D. Ramanan, C. L. Zitnick, and P. Dollár. Microsoft coco: Common objects in context, 2015.

G. W. Lindsay. Convolutional neural networks as a model of the visual system: Past, present, and future. Journal of cognitive neuroscience, 33(10):2017–2031, 2021.

T. Macpherson, A. Churchland, T. Sejnowski, J. DiCarlo, Y. Kamitani, H. Takahashi, and T. Hikida. Natural and artificial intelligence: A brief introduction to the interplay between ai and neuroscience research. Neural Networks, 144:603–613, 2021.

L. McIntosh, N. Maheswaranathan, A. Nayebi, S. Ganguli, and S. Baccus. Deep learning models of the retinal response to natural scenes. Advances in neural information processing systems, 29:1369–1377, 2016.

V. B. Mountcastle. Perceptual neuroscience: The cerebral cortex. Harvard University Press, 1998.

T. Naselaris, K. N. Kay, S. Nishimoto, and J. L. Gallant. Encoding and decoding in fMRI. NeuroImage, 56(2):400–410, 2011. ISSN 1053-8119.

D. Pitcher and L. G. Ungerleider. Evidence for a third visual pathway specialized for social perception. Trends in Cognitive Sciences, 25(2):100–110, 2021.

P. Poirazi, T. Brannon, and B. W. Mel. Pyramidal neuron as two-layer neural network. Neuron, 37(6):989–999, 2003.

R. Prenger, M. C.-K. Wu, S. V. David, and J. L. Gallant. Nonlinear v1 responses to natural scenes revealed by neural network analysis. Neural Networks, 17(5-6):663–679, 2004.

B. A. Richards, T. P. Lillicrap, P. Beaudoin, Y. Bengio, R. Bogacz, A. Christensen, C. Clopath, R. P. Costa, A. de Berker, S. Ganguli, et al. A deep learning framework for neuroscience. Nature neuroscience, 22(11):1761–1770, 2019.

M. Riesenhuber and T. Poggio. Computational models of object recognition in cortex: A review. 2000.

A. W. Roe, L. Chelazzi, C. E. Connor, B. R. Conway, I. Fujita, J. L. Gallant, H. Lu, and W. Vanduffel. Toward a unified theory of visual area v4. Neuron, 74(1):12–29, 2012.

P. H. Schiller and N. K. Logothetis. The color-opponent and broad-band channels of the primate visual system. Trends in neurosciences, 13(10):392–398, 1990.

M. T. Schmolesky, Y. Wang, D. P. Hanes, K. G. Thompson, S. Leutgeb, J. D. Schall, and A. G. Leventhal. Signal timing across the macaque visual system. Journal of neurophysiology, 79 (6):3272–3278, 1998.

K. Seeliger, L. Ambrogioni, Y. Güçlütürk, L. M. van den Bulk, U. Güçlü, and M. van Gerven. End-to-end neural system identification with neural information flow. PLOS Computational Biology, 17(2):e1008558, 2021.

G. St-Yves and T. Naselaris. The feature-weighted receptive field: an interpretable encoding model for complex feature spaces. NeuroImage, 180:188–202, 2018. ISSN 1053-8119. New advances in encoding and decoding of brain signals.

K. R. Storrs, T. C. Kietzmann, A. Walther, J. Mehrer, and N. Kriegeskorte. Diverse Deep Neural Networks All Predict Human Inferior Temporal Cortex Well, After Training and Fitting. Journal of Cognitive Neuroscience, 33(10):2044–2064, 09 2021. ISSN 0898-929X. doi: 10.1162/jocn_a_01755.

L. G. Ungerleider. Two cortical visual systems. Analysis of visual behavior, pages 549–586, 1982.

B. A. Wandell and J. Winawer. Imaging retinotopic maps in the human brain. Vision Research, 51(7):718–737, 2011. ISSN 0042-6989. Vision Research 50th Anniversary Issue: Part 1.

Y. Xu and M. Vaziri-Pashkam. Limits to visual representational correspondence between convolutional neural networks and the human brain. Nature communications, 12(1):1–16, 2021.

D. L. Yamins and J. J. DiCarlo. Using goal-driven deep learning models to understand sensory cortex. Nature neuroscience, 19(3):356–365, 2016.

D. L. K. Yamins, H. Hong, C. F. Cadieu, E. A. Solomon, D. Seibert, and J. J. DiCarlo. Performance-optimized hierarchical models predict neural responses in higher visual cortex. Proceedings of the National Academy of Sciences of the United States of America, 111 (23):8619–24, 2014. ISSN 1091-6490. doi: 10.1073/pnas.1403112111.

A. R. Zamir, A. Sax, W. Shen, L. J. Guibas, J. Malik, and S. Savarese. Taskonomy: Disentangling task transfer learning. In Proceedings of the IEEE conference on computer vision and pattern recognition, pages 3712–3722, 2018.

Y. Zhang, T. S. Lee, M. Li, F. Liu, and S. Tang. Convolutional neural network models of v1 responses to complex patterns. Journal of computational neuroscience, 46(1):33–54, 2019.

C. Zhuang, S. Yan, A. Nayebi, M. Schrimpf, M. C. Frank, J. J. DiCarlo, and D. L. Yamins. Un-supervised neural network models of the ventral visual stream. Proceedings of the National Academy of Sciences, 118(3), 2021.

