## Supplementary Information for "Brain-optimized neural networks learn non-hierarchical models of representation in human visual cortex"

#### Supplementary tables

| Abbreviation | Description |
|---|---|
| Gabor-gpf | Gabor wavelet feature extractor with a single voxelwise Gaussian pooling field. |
| AlexNet-gpf | AlexNet feature extractor with a single voxelwise Gaussian pooling field. |
| AlexNet-fpf | AlexNet feature extractor with one flexible pooling field for each feature map resolution. |
| GNet8j-fpf | GNet feature extractor trained on all brain area jointly (j) on 8 subjects, with one flexible pooling field for each feature map resolution. |
| GNet8jft-fpf | GNet8j-fpf model with fine tuning optimization procedure. |
| GNet8r-fpf | GNet feature extractor trained each brain area separately, i.e. ROI-wise (r), on 8 subjects, with one flexible pooling field for each feature map resolution. |

**Supplementary Table 1: Table of models.** A brief description of the various encoding models.

| Symbol | Description |
|---|---|
| $x$ | An image |
| $r$ | Empirical voxel activity |
| $\bar{r}$ | Voxel prediction |
| $\bar{r}_v$ | Voxel prediction of voxel $v$ |
| $V$ | A population of voxels |
| $\bar{r}^{\text{model}}$ | Voxel prediction of a specific model |
| $\rho^{\text{model}}$ | Pearson correlation between prediction of model and target activity (context dependent) |
| $e_l(x)$ | $l$-th set of feature map of $x$ |
| $[e_l(x)]_{kji}$ | A specific feature of feature map $e_l(x)$ |
| $\Phi(x)$ | Spatially pooled feature of $x$ |
| $g_v^n$ | $n$-th pooling field of voxel $v$ |
| $w_v$ | Feature weight for voxel $v$ |
| $\langle\cdot\rangle_V$ | Average over voxel population $V$ |

**Supplementary Table 2: Table of symbols** Description of the most frequently used symbols.

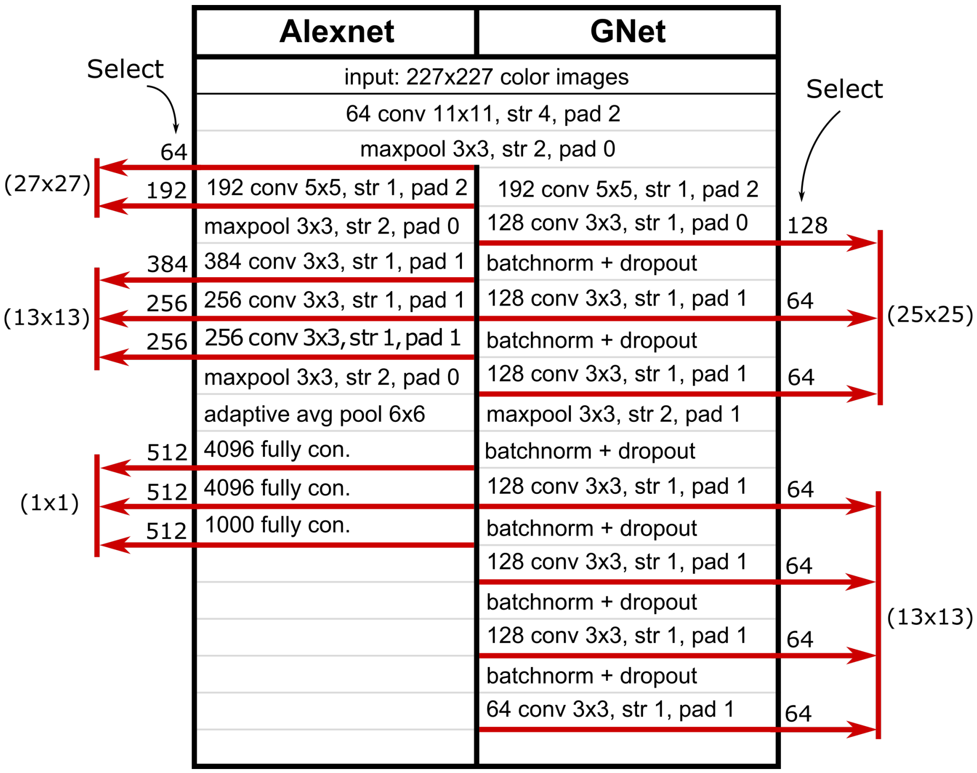


**Supplementary Table 3: Details of the feature extractor networks**. Layer-by-layer network structure comparison between AlexNet and GNet. The number of feature maps connected to the read-out heads is indicated under the 'select' label. In the case of AlexNet, this selection is based on feature map variance w.r.t the NSD training set, whereas the GNet selection is based on a fixed partition of the feature maps at onset of training.

### Supplementary figures

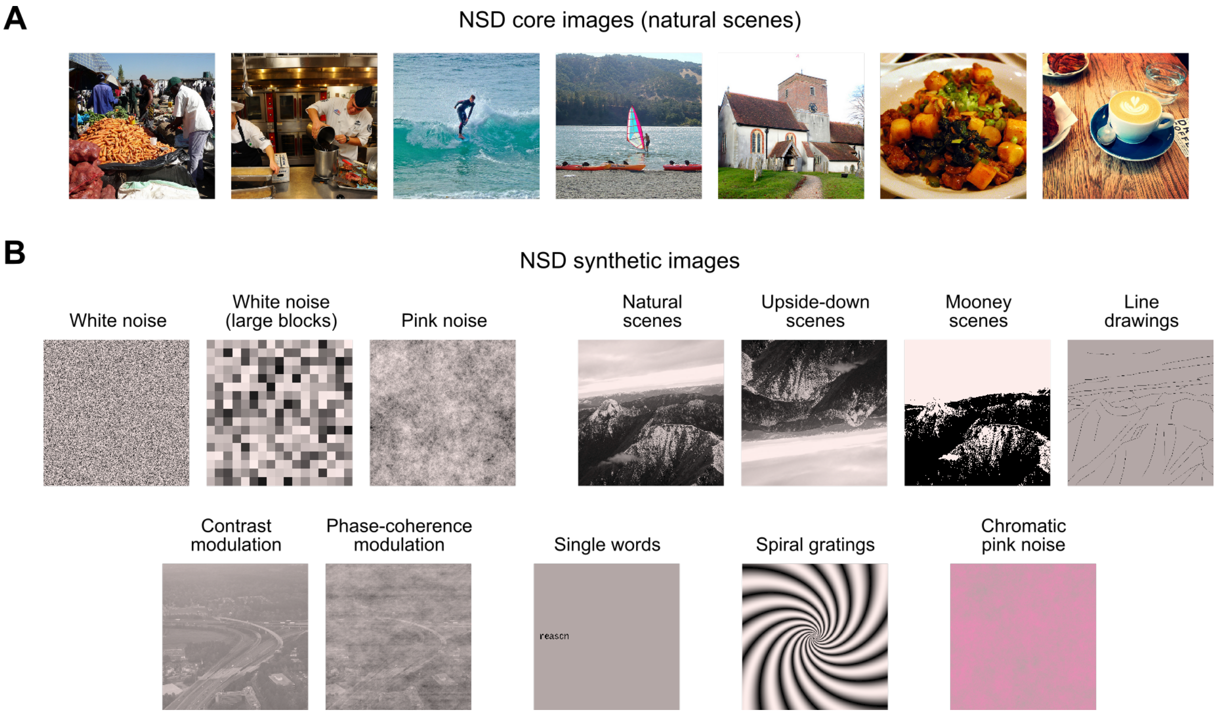


**Supplementary Figure 1: Examples of NSD stimuli.** A) Example images from the core NSD experiment. The images are natural scenes pulled from the Microsoft Common Objects in Context database. B) Example images from the NSD synthetic experiment. The images follow certain conceptual groupings, as depicted (see Methods for details).
